# Genome-wide sampling suggests island divergence accompanied by cryptic epigenetic plasticity in Canada lynx

**DOI:** 10.1101/316711

**Authors:** J.B. Johnson, D.L. Murray, A.B.A. Shafer

**Affiliations:** Environmental and Life Sciences Graduate Program, Trent University, K9J 7B8 Peterborough, Canada; Biology Department, Trent University, K9J 7B8 Peterborough, Canada; Forensics Department, Trent University, K9J 7B8 Peterborough, Canada

**Author notes:** **Corresponding author**: J.B. Johnson, Address: 1600 W Bank Drive. Peterborough, ON, Canada, K9J 7B8. https://orcid.org/0000-0001-5077-4096.

**Keywords:** ecological epigenetics, population differentiation, island biogeography

## Abstract

Determining the molecular signatures of adaptive differentiation is a fundamental component of evolutionary biology. A key challenge remains for identifying such signatures in wild organisms, particularly between populations of highly mobile species that undergo substantial gene flow. The Canada lynx (*Lynx canadensis*) is one species where mainland populations appear largely undifferentiated at traditional genetic markers, despite inhabiting diverse environments and displaying phenotypic variation. Here, we used high-throughput sequencing to investigate both neutral genetic structure and epigenetic differentiation across the distributional range of Canada lynx. Using a customized bioinformatics pipeline we scored both neutral SNPs and methylated nucleotides across the lynx genome. Newfoundland lynx were identified as the most differentiated population at neutral genetic markers, with diffusion approximations of allele frequencies indicating that divergence from the panmictic mainland occurred at the end of the last glaciation, with minimal contemporary admixture. In contrast, epigenetic structure revealed hidden levels of differentiation across the range coincident with environmental determinants including winter conditions, particularly in the peripheral Newfoundland and Alaskan populations. Several biological pathways related to morphology were differentially methylated between populations, with Newfoundland being disproportionately methylated for genes that could explain the observed island dwarfism. Our results indicate that epigenetic modifications, specifically DNA methylation, are powerful markers to investigate population differentiation and functional plasticity in wild and non-model systems.

**SIGNIFICANCE:** Populations experiencing high rates of gene flow often appear undifferentiated at neutral genetic markers, despite often extensive environmental and phenotypic variation. We examined genome-wide genetic differentiation and DNA methylation between three interconnected regions and one insular population of Canada lynx (*Lynx canadensis)* to determine if epigenetic modifications characterized climatic associations and functional molecular plasticity. Demographic approximations indicated divergence of Newfoundland during the last glaciation, while cryptic epigenetic structure identified putatively functional differentiation that might explain island dwarfism. Our study suggests that DNA methylation is a useful marker for differentiating wild populations, particularly when faced with functional plasticity and low genetic differentiation.

## INTRODUCTION

Investigations into adaptive differentiation have often relied on either quantifying phenotypic variation or contrasting genetic polymorphisms identified in an organism’s DNA (1, 2). Environmental conditions can be a powerful driver of population differentiation and phenotypic plasticity, where relationships are traditionally ascertained by correlating allele frequencies to environmental and morphological variation (3). Detecting adaptive differentiation in species that experience high rates of gene flow, however, is challenging due to the homogenization of genomic regions that are neutral or under weak selection (4). An aspect of molecular variation that is undetected by standard genetic sequencing involves direct modifications to the structure of DNA. Epigenetic modifications like DNA methylation are influenced by environmental conditions, directly affect gene expression, and may even be indicative of early divergence due to local adaptation (5–7).

DNA methylation has been implicated in mediating ecologically relevant traits due to varied environmental conditions (8), primarily due to its regulatory role in transcription by modifying chromatin structure, repressing transcription factors, or recruiting protein complexes that block transcriptional machinery, especially around CpG islands (9–11). CpG islands are dense clusters of cytosine-guanine dinucleotides and frequently occur near the transcription start site of genes and have a functional relationship with gene expression (12–14). Consequently, DNA methylation, particularly around genes that impart an ecologically relevant benefit, could be used to characterize populations that exhibit low levels of neutral genetic differentiation despite extensive differences in resources and conditions.

Here, we assessed whether environmental variation, geographic distance, or insularity were determinants of DNA methylation structure in a free-ranging carnivore. Our study species, the Canada lynx (*Lynx canadensis*), is a mid-sized felid that is highly mobile and whose neutral genetic variation (i.e., microsatellites) exhibits low levels of genetic differentiation across the mainland, with divergent island populations (15, 16). Although some hypotheses exist regarding the colonization of Newfoundland (15), no formal demographic analyses have been investigated using genome-wide markers. Despite a low degree of overall genetic structure, we hypothesize that two potential mechanisms might drive epigenetic divergence in Canada lynx. First, allele frequencies are correlated to climatic gradients in both population time-series and fine-scale genetic analyses (15, 17, 18), suggesting climate might influence patterns of methylation in lynx due to its semi-plastic nature. Second, Canada lynx show a subtle cline in body size with larger individuals in Alaska (19) to smaller individuals in insular populations including Newfoundland and Cape Breton Island (20). Body size changes in island populations appear to be consistent with the ‘Island Rule’ (20), where insular mammals are smaller in size compared to their mainland counterparts (21–24). If functional genes related to body size are repressed in geographically isolated populations, then epigenetic modifications might underlie the molecular plasticity that is associated with these differences (25). Based on these two mechanisms (climate and island isolation), we tested two predictions: i) climatic conditions are the driving force behind spatial epigenetic structuring (26); and ii) Canada lynx on the island of Newfoundland exhibit differential methylation compared to mainland populations that explains the observed trends in phenotypic divergence (27).

## RESULTS

We collected 95 Canada lynx epidermal tissue samples from four populations (n = 23-24 per sampling area) across North America, including one insular population in Newfoundland (Fig. 1; Table S1. The sampled populations have a wide geographical spread, with an average minimum distance between populations (Québec and Newfoundland) of 1,158 km and a maximum distance (Alaska and Newfoundland) of 5,520 km. Habitats around these populations present a dynamic range of environmental conditions, ranging from 32 to 432 mm of winter precipitation and a mean annual temperature range (28) of −6.3 to 4.7° C. To determine associations between these environmental conditions and genome-wide patterns of DNA methylation, we created a reduced representation bisulfite sequencing (RRBS) library (full protocol in Supplemental). Paired-end sequencing on a HiSeq2500 generated a total of 210,773,612 filtered and demultiplexed reads that were aligned to the domestic cat genome (85.0% average mapping success; Table S1 *Felis catus*; NCBI: felcat9.0) and variants were called using specially designed software for bisulfite converted reads (29). The cat genome was chosen due to our interests in functional annotations and the multiple revisions this genome has underwent resulting in a chromosome-scale assembly (NCBI: felcat9.0; 3GCA_000181335.4). We assessed bisulfite conversion efficiency by including non-methylated lambda phage DNA in the sequencing lane, and mapped our reads to the human (*Homo sapiens*) and lambda phage genomes to rule out contamination (2.7% and <0.1% success, respectively). We used a Bayesian framework to call genetic variants from bisulfite converted reads; these SNPs were used to characterize historical demography and neutral genetic structure. The distribution of methylated sites and SNPs are shown in Fig. 2; our multiplexed RRBS library with 95 individuals sampled an average of 2.6 million unique CpG positions per individual (*Min* = 262,000; *Max* = 9,677,599; Table S2). Furthermore, we quantified the temporal effects of methylation, the ramifications of missing data, model parameterization, and the effects of dataset subdivisions based on feature type with sensitivity analyses, with no implications on overall inferences (Fig. S1 – S5).

**Fig. 1.**
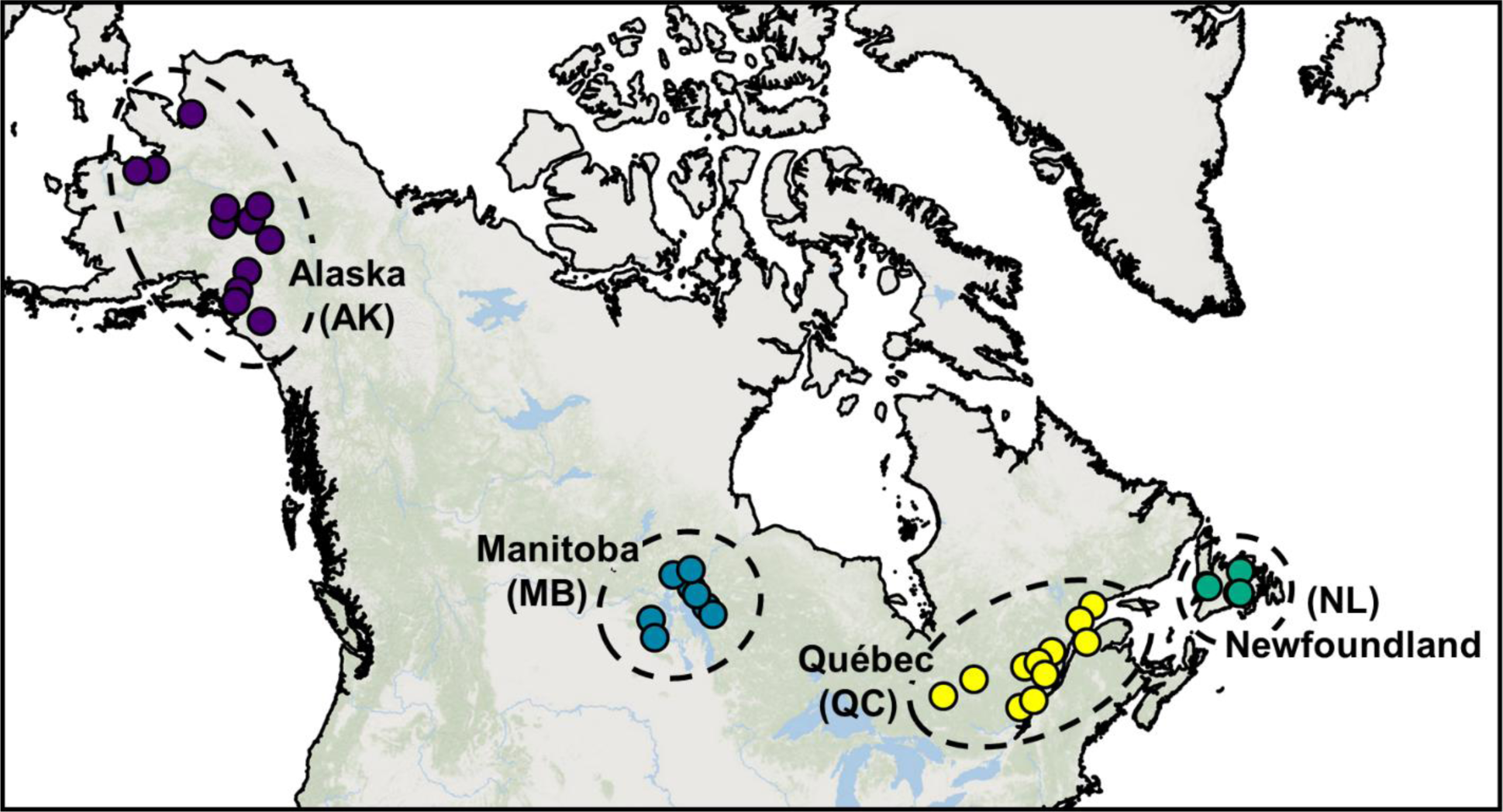
Distribution of 95 Canada lynx (*Lynx canadensis*) samples across North America, subjected to bisulfite conversion to assess site specific differentially methylated variations between populations. All four populations are delineated by colour and include 24 individuals, except Alaska (n = 23).

**Fig. 2.**
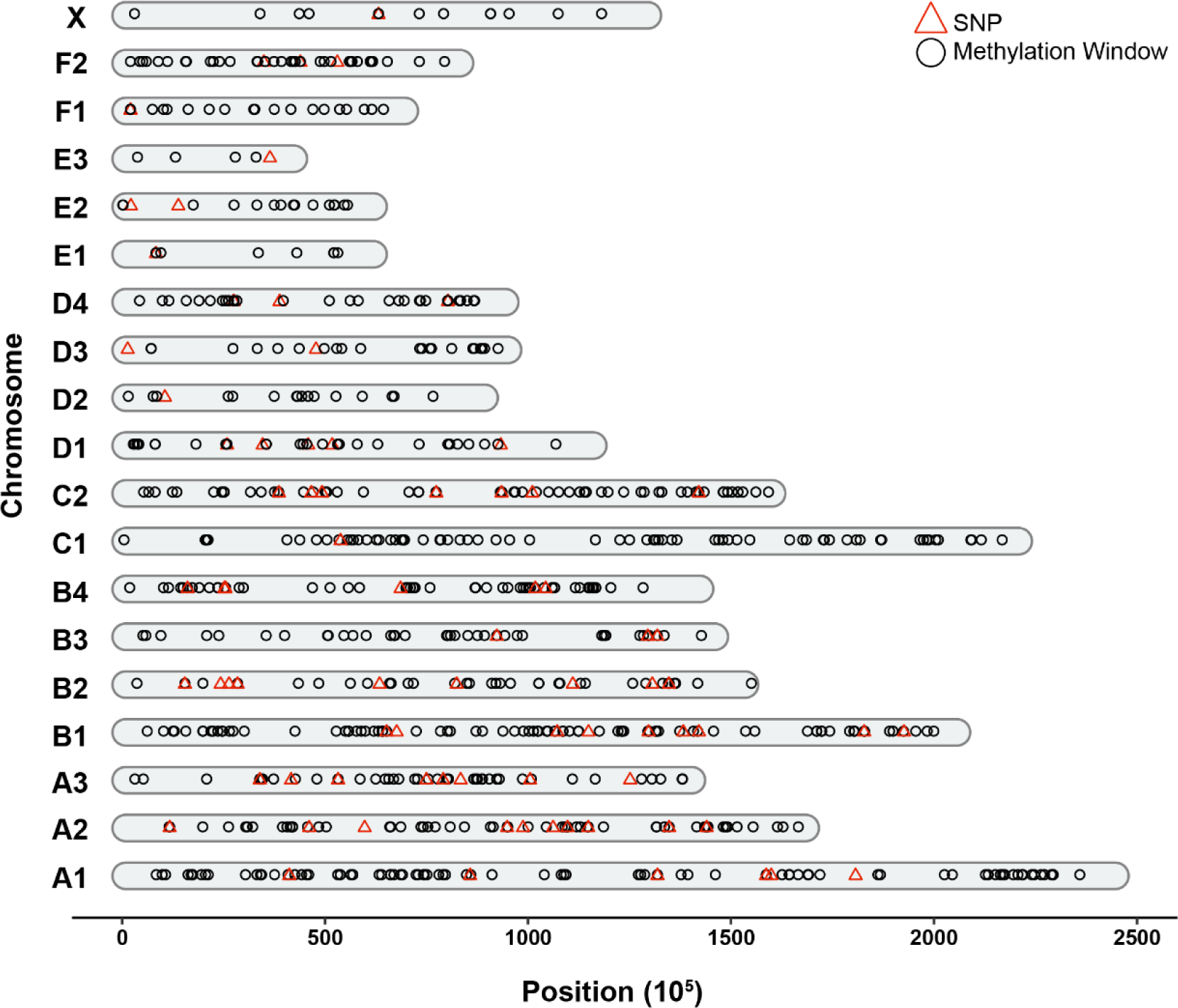
Sampling of the genome in this reduced-representation bisulfite sequencing study for 95 Canada lynx, where filtered reads were mapped to the 19 chromosomes of the domestic cat.

### Population demography and subtle neutral genetic differentiation

We estimated the time of divergence (*τ*_*i*_) and the best fit historical demographic model for Newfoundland using an AIC model selection framework using the SNP dataset. Demographic scenarios (Table S3) were determined using the site-frequency spectrum (SFS) implemented in the program ∂a∂i (30). The best fit model for our joint SFS (Fig. S6) between the mainland and Newfoundland suggests a split during the last glacial maximum (*τ*_*i*_ = 26.5 ± 6.5 *KYA*, Table S4) with early asymmetric admixture from the mainland into the ancestral Newfoundland population (Fig. 3). This model indicated that Newfoundland likely experienced a founder effect with no contemporary admixture with the mainland, while overall diversity between populations was still relatively low (θ = 3.1). An additional one-dimensional SFS solely for Newfoundland corroborated our identification of a demographic bottleneck (Fig. S7). We then performed an analysis of molecular variance (AMOVA) to assess relative population differentiation, where extensive variation was seen within samples (*σ* = 24.8) relative to overall variation between populations (*σ* = 0.46). Relative differences among all populations (*ϕ* = 0.02) was again diminutive compared to the variation determined between Newfoundland and the mainland (*ϕ* = 0.09). Bayescan identified no highly differentiated (*F*_*ST*_) loci under selection (Fig. S8), and a general pattern consistent with isolation by distance was observed (*pseudo* - ***F*** = 43.6; *R*^2^ = 0.31; *p* ≤ 0.001). A principal coordinates analysis computed from a Euclidean distance matrix summarizing SNP variation detected relatively congruous populations (Fig. 4), where the first axis largely described variation between the mainland and Newfoundland (*PCo*1 = 30.3%), while the second axis detected the subtler differentiation between mainland populations (*PCo*2 = 6.1%).

**Fig. 3.**
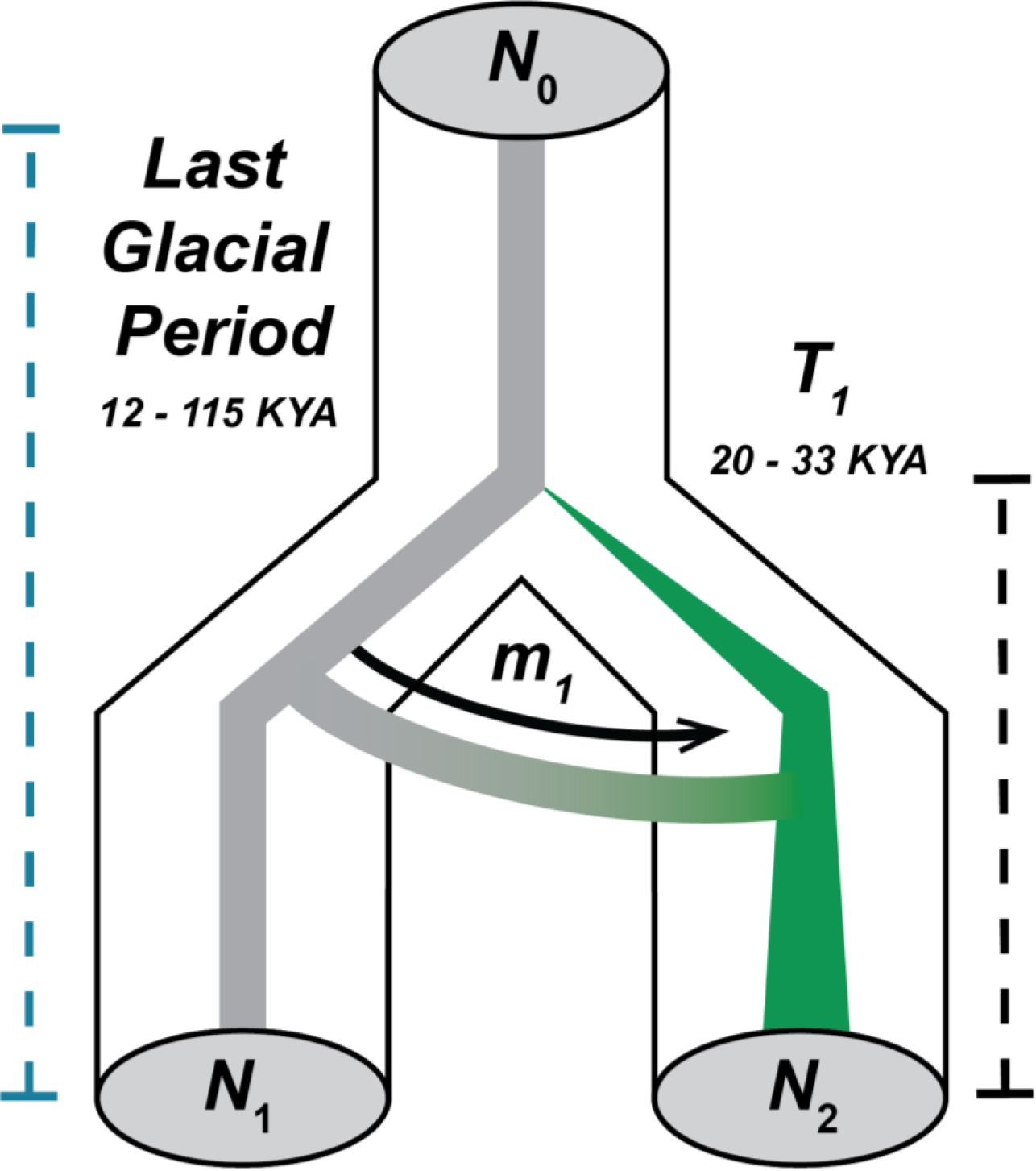
Schematic of the most likely demographic trajectory of mainland (N_1_) and Newfoundland (N_2_) Canada lynx, as identified by the lowest AIC during model selection using the site-frequency spectrum (SFS) determined using ∂a∂i. The model suggests divergence with early asymmetric migration into the Newfoundland population. The Newfoundland population exhibits signatures of a founder effect, with a recent increase in N_e_ and a lack of gene flow with the mainland.

**Fig. 4.**
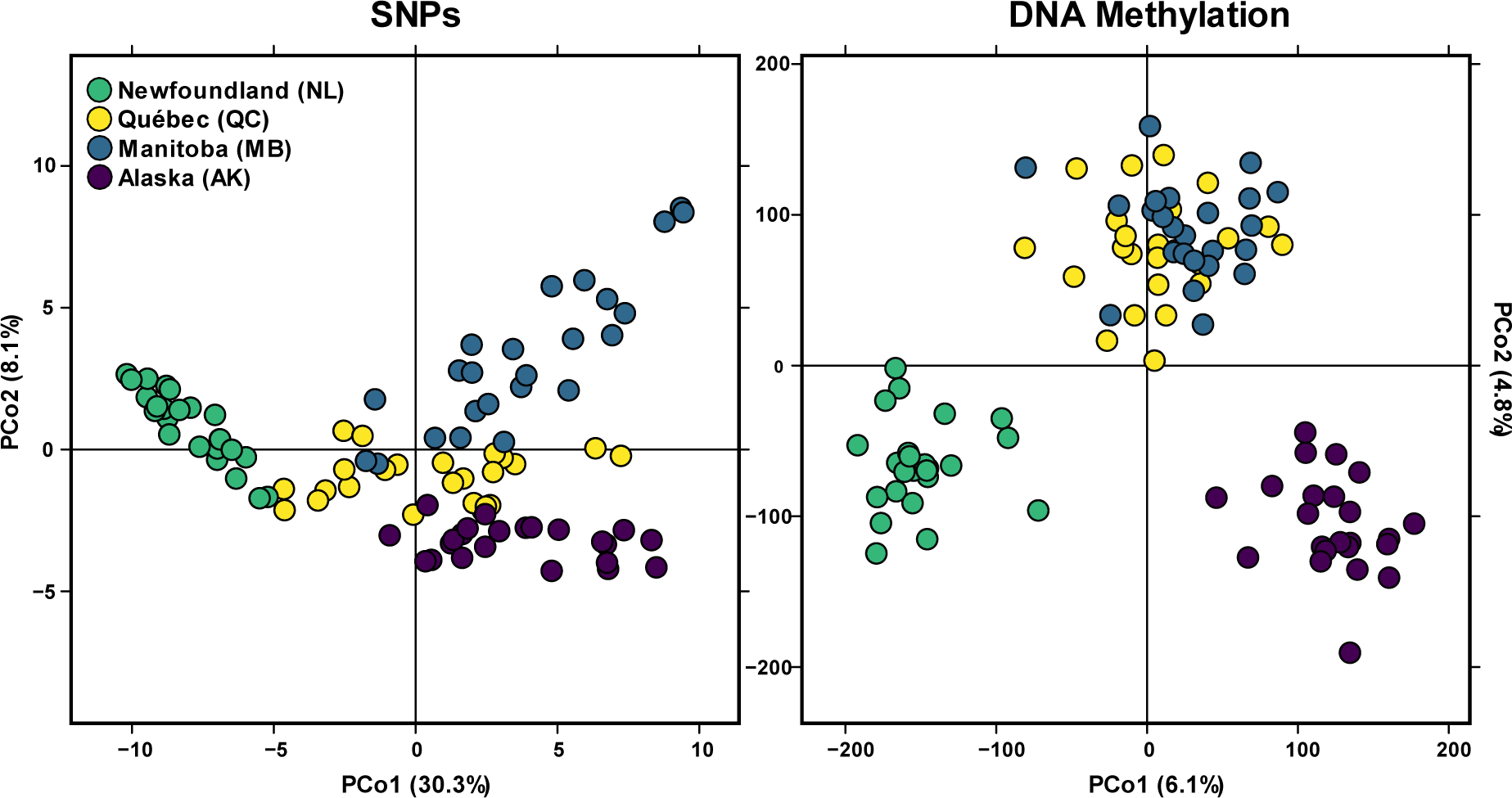
Principal coordinate ordination plots of genetic (left) and epigenetic (right) variation between Canada lynx populations indicate cryptic structure at epigenetic markers, with individuals as points and populations delineated by colour. All molecular data was summarized with a pair-wise Euclidean dissimilarity matrix. Methylation data was summarized with 5,000-bp running windows across the genome (n = 705) and SNPs were called from bisulfite converted reads (n = 85).

### Pronounced population structure revealed by methylation patterns

DNA methylation patterns were examined across the genome (Fig. 2) in 5,000-bp windows, resulting in a total of 705 windows with 9,642 CpGs for analyses after filtering for coverage and equal minimum representation across all populations. A principal coordinates analysis (PCoA) summarizing methylation variation (*PCo*1 = 6.1%, *PCo*2 = 4.8% variation explained) showed slightly disparate trends to the SNP dataset of neutral population structure, where variation in DNA methylation clustered Alaska away from the remaining mainland populations and indicated no subdivisions between the mid-continental populations (Fig. 4). The first PCoA axis roughly indicated a pattern of isolation by distance, while the second axis identified unique variation separating the geographically peripheral populations (Newfoundland and Alaska) from the mid-continental populations.

We used multivariate distance-based redundancy analyses (db-RDAs) and step-wise model selection to quantify the determinants of observed variation in DNA methylation, which identified three significant variables: geographic distance, winter conditions and a binary variable representing insularity for the Newfoundland population (*pseudo* - ***F*** = 16.6; *R*^2^ = 0.35; all variables *p* ≤ 0.001, Table S5). Tree cover (*p* = 0.77) and a randomly-generated numerical variable to assess the effect of noise (*p* = 0.71) added no explanatory power to the model (Table S5). Collinearity was low between all retained variables (*VIF* = 2.09 - 3.77; Table S6). We examined the explanatory power of each variable independently using partial db-RDAs (Fig. 5), which identified the most variation solely explained by geographic distance (*pseudo* - ***F*** = 17.1; *R*^2^ = 0.12; *p* ≤ 0.001), the least explained by winter conditions (*pseudo* - ***F*** = 7.1; *R*^2^ = 0.05; *p* ≤ 0.001), anqed an intermediate amount explained by the largely impassable barrier of the Strait of Belle Isle (*pseudo* - ***F*** = 13.5; *R*^2^ = 0.10; *p* ≤ 0.001). Initial analyses were conducted to determine if *a priori* genomic feature-type classification would affect downstream conclusions.

**Fig. 5.**
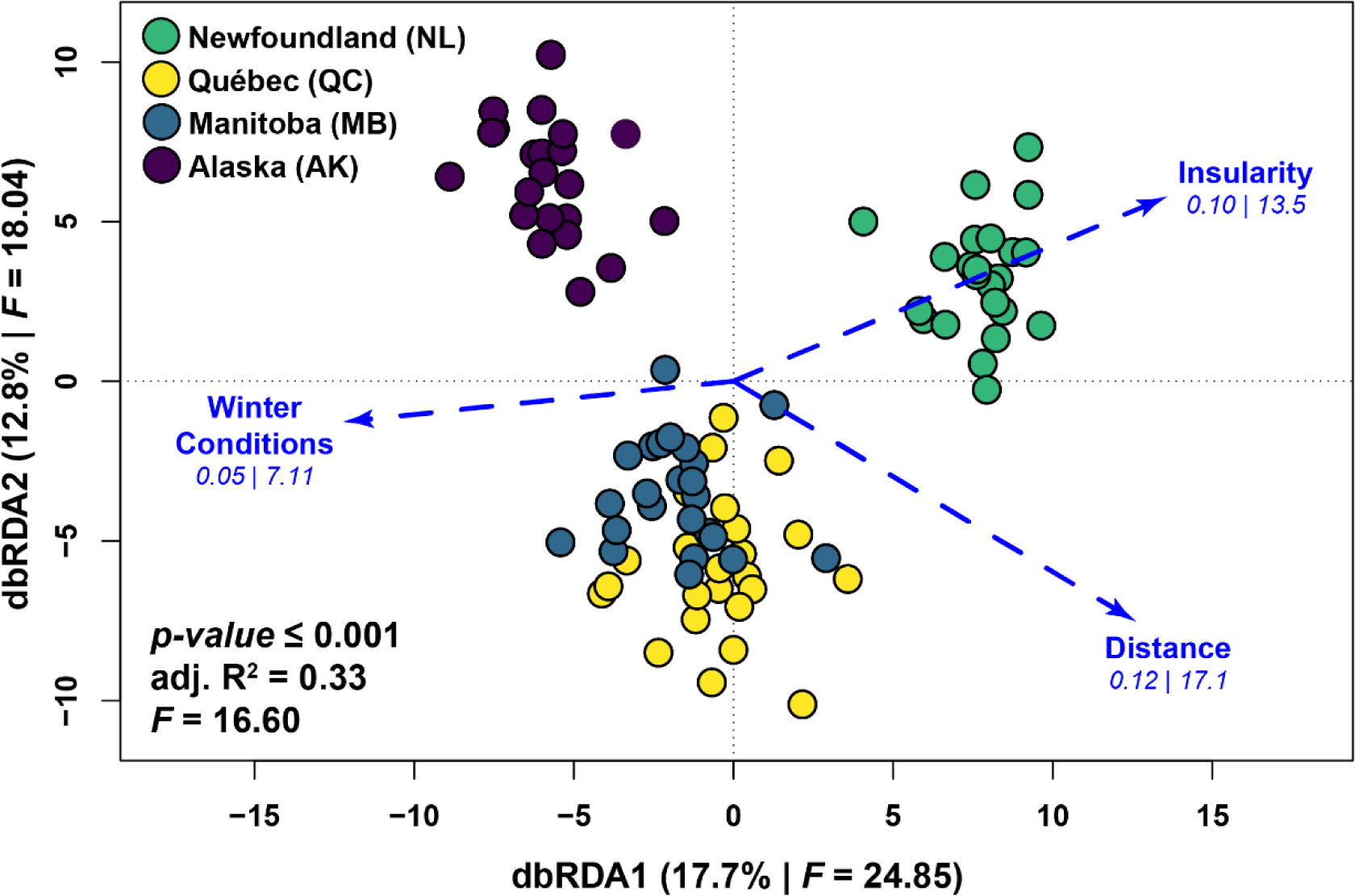
Distance-based redundancy analysis (db-RDA) ordination on DNA methylation indicate geographic distance, insularity, and winter conditions explain a portion of the observed epigenetic variation, where populations are delineated by colour. The axes of a principal coordinates analysis (PCoA) summarizing methylation data were used a response variable to determine biogeographical relationships. Values underneath each environmental vector indicate R^2^ and ***F*** values obtained from partial db-RDAs.

### DMRs over morphological genes could explain island dwarfism

We conducted a *de novo* annotation of CpG islands throughout the cat genome (*N* = 28,127) using hidden Markov models, which contained an average GC content of 59.2% with a posterior probability of observed-to-expected GC content (CpG_o/e_) of 1.13. Our 705 5,000-bp windows of methylation were then divided based on overlap with a CpG island or gene body, which were used to identify differential methylation between populations using beta regressions with a Bonferroni-corrected significance threshold of 3.19 × 10^-5^. A total of 42 differentially methylated genes were identified across all populations, including 4 long non-coding RNAs. Overrepresented functional pathways for differentially methylated genes were identified using a Panther-Gene Ontology (GO) analysis, which identified cellular component morphogenesis (GO: 0032989, *n* = 4, *p* = 0.007) and embryonic development (GO: 0009790, *n* = 3, *p* ≤ 0.001) as the most overrepresented biological pathways (Table S7). Three specific genes identified with the ontology analysis warranted closer examination due to their known direct regulation of body mass and morphology, as identified in previous laboratory experiments. These genes (HDAC9, TMOD2, and ZEB1) were hypermethylated in Newfoundland compared to the mainland, with a difference in methylation of 39.6, 29.4, and 37.6%, respectively (Fig. 6).

**Fig. 6.**
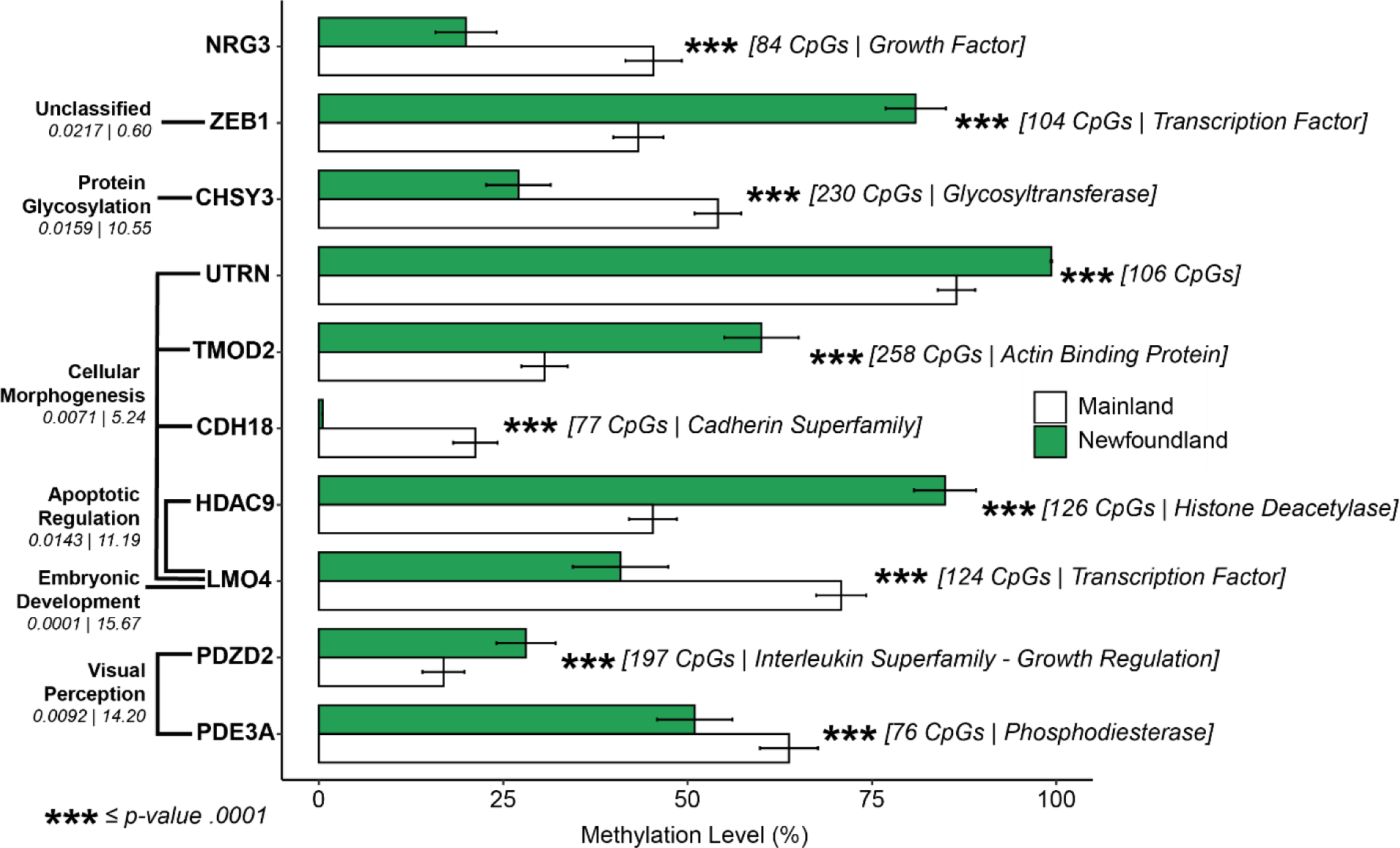
Enriched differentially methylated genes, highlighting differences between Newfoundland and Mainland Canada lynx. Methylation levels are indicated with standard error of the mean, with all mainland populations lumped together. Enriched biological processes were identified using a GO-Panther analysis, with *p-value* and enrichment values indicated under each process (far left). The total number of CpGs per gene analyzed across populations and the GO-Panther gene-specific function is identified to the right of the bar plots. All differentially methylated regions were located directly over gene bodies.

## DISCUSSION

### Newfoundland as a glacial refuge for Canada lynx

The historical demography of insular Canada lynx has been the focus of speculation, with estimates ranging from intra-glacial colonization to introductions within the last century (15). Our demographic analyses with SNP data suggest the former, where colonization occurred sometime during the last-glacial period; for example an ice bridge is believed to have connected the mainland to Newfoundland and portions of the island were still thought to be ice-free (31). While our population divergence time estimates are scaled by mutation rate and generation time, all reasonable metrics support a historic colonization (> 1,000 years). Our analyses also indicated that Newfoundland underwent a demographic bottleneck after colonization, consistent with microsatellite data (16), and that genetic diversity remains low compared to mainland populations. Row et al. (15) showed a minimal barrier effect of the Rocky Mountains to Canada lynx, which is also supported with our SNP data. Despite minimal range-wide nuclear divergence in lynx, it remains possible that distinct environmental variation might be driving unique patterns of functionally important differentiation at the range margins, including Alaska and Newfoundland.

### DNA methylation detects biologically relevant cryptic population structure

We have shown that despite largely undifferentiated populations across the mainland at the genetic level, epigenetic structure across the range is substantial and is suggestive of functional epigenetic plasticity in the face of genetic homogeneity. Despite the lack of genetic structure between mainland populations (Rueness et al. 2003, this study), we observed distinct patterns of DNA methylation that differentiate Alaska from mid- and east-continental populations. While the segregation of Newfoundland from the mainland was largely reflected across both epigenetic and genetic datasets, closer examination of differentially methylated regions in Newfoundland appear to explain the phenotypic trends observed in this insular population. Specifically, the differentiated DNA methylation patterns in the Newfoundland population suggest a molecular pathway for the morphological differences observed between mainland and island populations (19, 20), as predicted by the Island Rule (24).

Hypermethylation and thus putative downregulation of genes directly implicated in reduced body mass (HDAC9, TMOD2, ZEB1) suggests that epigenetic modifications could be an underlying mechanism in island dwarfism (33–36). Genetic knockout of HDAC9 in laboratory experiments with mice (*Mus musculus)* have indicated diminished body mass (33), whereas hypermethylation in the Newfoundland population suggests an analogous trend of repressed gene expression that could explain the diminished size of these lynx. Similarly, TMOD2 is a member of the tropomodulins family, which are associated with actin regulation where downregulation associates with a decrease in cell height during morphogenesis (34, 35). Hypermethylation of the zinc finger transcription factor ZEB1 in the Newfoundland population also provides evidence for the role of DNA methylation in island dwarfism. High levels of ZEB1 are associated with larger body mass in mice (36), whereas hypermethylation in Newfoundland suggests reduced expression of ZEB1 and again associates with the smaller size of insular lynx relative to their mainland counterparts (20).

The genetic contribution to epigenetic variation cannot be ignored (37), but cannot be irrefutably resolved without an economically-prohibitive amount of sequencing, including whole-genome and whole-methylome assays (6). Our genome-wide sampling of a non-model organism and localization of SNPs completely outside of DMRs indicates that these genetic data largely describe neutral genetic structure without confounding our signals from methylation data, while our epigenetic data describe molecular plasticity with potential functional ramifications.

Importantly, our demographic inferences suggesting a relatively recent evolutionary establishment of Newfoundland Canada lynx could implicate DNA methylation as a marker to examine rapid evolutionary change. Evolutionary theory in model organisms has predicted that epimutations and methylome evolution often precede genomic changes (38), which has been further substantiated with empirical evidence examining epigenetic changes over structural genomic variants in the absence of genetic variation (39).

As evidence accumulates regarding the mechanisms underlying epigenetic inheritance (40) and the adaptive benefits of epialleles (41), DNA methylation has the potential to be worked into a framework of adaptive divergence (42, 43) and ultimately into the ecological speciation model (4). It is possible that hypermethylation of locally deleterious genes could provide a precursory mechanism to adaptive divergence over linked SNPs. In contrast, if phenotypically-plastic epigenetic marks are advantageous and heritable, then selection might act directly on the repression of these genomic locations, even in the absence of underlying genetic divergence, particularly if such genes are metabolically costly and provide no fitness benefit, if activated (44). Provided that the mechanisms underlying transgenerational epigenetic inheritance can maintain methylomic patterning across generations (Colome-Tatche et al. 2012, Schmitz et al. 2013, Liu et al. 2018, Joo et al. 2018, but see Kazachenka et al. 2018), DNA methylation could be integrated into the working model of ecological speciation with gene flow by mediating gene expression and the optimal phenotype under local conditions.

Overall, our results indicate that DNA methylation can detect cryptic population structure in genetically homogenous species while providing insight on the functional pathways that shape phenotypic divergence between populations. Epigenetic modifications offer great utility for understanding the mechanisms intertwining ecology and evolution (50, 51), and we offer among the first empirical results demonstrating the historical and functional inferences that can be gained from comparative methylomics, even in non-model organisms.

## METHODS

### Sample Acquisition and Reduced-Representation Bisulfite Sequencing (RRBS) Library Preparation

Georeferenced Canada lynx tissue samples were collected from fur auction houses throughout eastern North America from dried pelts (North American Fur Auctions, Fur Harvester’s Auctions, Inc.). Individuals from four geographic locations were chosen for this study, spanning the longitudinal and latitudinal range of the species and consistent with previous research on the Canada lynx system (32). Tissue was consistently taken from the same morphological location from each adult-sized pelt, although sex could not be determined. Genomic DNA was isolated using a MagneSil^®^ (Promega Corporation) Blood Genomic Max Yield System a JANUS^®^ workstation (PerkinElmer, Inc.) and quantified and standardized to 20 ng/μl with a Quant-iT PicoGreen^®^ ds-DNA assay using manufacturer’s instructions (Thermo Fisher Scientific). We adapted an existing reduced representation bisulfite sequencing library preparation workflow designed for multiplexed high-throughput sequencing (52). Genomic DNA (400 ng) for 95 Canada lynx samples was digested with NsiI and AseI restriction enzymes overnight and subsequently ligated with methylated adapters (Table S8). An individual sample of completely non-methylated lambda phage genomic DNA (200 ng; Sigma-Aldrich – D3654) with a unique barcode was included to assess bisulfite conversion efficiency (Table S9). Barcoded samples were then combined into eight pools to ensure consistent reaction environments for the entire library using a QIAquick PCR Purification Kit (Qiagen, Valencia, CA, USA) following manufacturer’s instructions and 10 μL of 3M NaAc was added to neutralize pH. We then performed a SPRI size selection on each pool (0.8x volume ratio) with Agencourt^®^ AMPure^®^ XP beads (Beckman Coulter, Inc.). Nicks between the 3’ fragment overhang and the 5’ non-phosphorylated adapter nucleotide were repaired with DNA polymerase I and bisulfite conversion was performed on each pool using an EZ DNA Methylation-Lightning Kit™ (Zymo Research) with a 20-minute desulphonation time.

Pools were amplified in three separate PCRs to mitigate stochastic differences in amplification and were subsequently concentrated using a QIAquick PCR Purification Kit (Qiagen, Valencia, CA, USA). The eight pools were quantified with Qubit 3.0^®^ (Thermo Fisher Scientific) and appropriate amounts were added to a final super-pool for equal weighting. A final magnetic bead clean-up (0.8x volume ratio) was performed to remove any adapter dimer. We checked final library concentration and fragment distribution with a Qubit^®^ 3.0 (ThermoFisher Scientific) and Hi-Sense Bioanalyzer 2100 chip (Agilent), respectively (Fig. S9). Paired-end 125-bp sequencing was performed on a single lane on an Illumina HiSeq 2500 platform with updated software appropriate for BS-seq datasets (HCS v2.2.68 / RTA v1.18.66.3) at the Centre for Applied Genomics at the Hospital for Sick Children (Toronto, Ontario, Canada), which has been determined to be one of the most reliable sequencing platforms for bisulfite-converted libraries (53). Raw sequence data FastQ files are available on the Sequence Read Archive (Accession: PRJNA509991). All bioinformatic and analytical code is available on GitLab (https://gitlab.com/WiDGeT_TrentU/RADseq/).

### Bioinformatics – quality checks and SNP calling

We assessed sequencing success as well as removed adapter and low-quality reads via FastQC (54) and Cutadapt (55) implemented in TrimGalore! v0.4.4 (56). Individuals were demultiplexed using python scripts (52). Paired-ends reads for Canada lynx samples were initially aligned to several genomes (*Felis catus, Homo sapiens,* Lambda Phage) using Bismark (29) to assess mapping efficiency and contamination (Table S1). Paired-end reads for downstream analyses were aligned to the domestic cat genome with Bismark, using Bowtie2 default settings and parameters seen in other studies (*score*_ min *L*, 0, −0.6) (29, 57). We mapped our reads to the cat genome, as opposed to the Iberian lynx (*L. pardinus,* Abascal et al. 2016) or the newly released Canada lynx (*L. canadensis*) genomes, due to the high-quality annotations and scaffolds of the cat genome, as we were interested specifically in functionally related regions and the cat genome has benefited from numerous revisions. While our choice of reference genome likely diminished total aligned reads, any undetermined downstream implications would affect all samples equally. We chose to map reads using Bismark due to its quantitative accuracy and speed at mapping a large number of samples (59).

SNPs were called from indexed BAM files with CGmapTools (60) using a coupled Bayesian wildcard algorithm with a conservative 0.01 error rate and a static 0.001 p-value for calling variant sites, which generated variant call files (VCFs). SNP calling from bisulfite-converted reads is amenable for defining population structure (61). VCF files were indexed, merged, and filtered using VCFtools (62) for bi-allelic loci with a sequencing depth of at least five, and were shared between at least 50% of the individuals (max-missing 0.5) and a minor allele frequency (maf) of > 0.001. Sites were further filtered by removing any variants out of Hardy-Weinberg equilibrium within populations (*p* < 0.05). A pair-wise Euclidean dissimilarity (distance) matrix was computed on the SNP data using the function *daisy* within the package *cluster* (63) using R v3.4.2 (64). This dissimilarity matrix was then summarized in a principal coordinates analysis (PCoA) using the *dudi.pco* function in *adegenet* (65). Missing data were imputed by mean allele at a population level. Pairwise *F*_*ST*_ was calculated using VCFtools (62) and any highly fixed SNPs putatively under selection were identified with BayeScan v2.0 with three runs using different prior odds ratios to test for sensitivity (66) (Fig. S8). AMOVA between populations was computed using the *poppr.amova* function within *poppr* (Excoffier et al. 1992, Kamvar et al. 2014, Fig. S10). We determined if genetic structuring was consistent with patterns of isolation by distance using a db-RDA with the first 3 axes of a PCoA on SNP data as response variables with the first axis of a PCoA on a Euclidean distance matrix of latitude and longitude as an explanatory variable.

### Demographic Inferences and Simulations

Historical demography was assessed using the program ∂a∂i (30). ∂a∂i performs numerous optimizations to assess fit between various demographic scenarios and the expected allele frequency spectrum according to a multi-population diffusion equation, which can then be repeated numerous times to find ideal optimization parameters and assess goodness of fit. VCF files were converted to a ∂a∂i appropriate SNP format using custom python scripts. Each locus contained a maximum of one SNP and thus assumptions of linkage disequilibrium for the SFS were satisfied.

To determine the best fit demographic model, two rounds of cross-validated model selection were conducted with 10 and 20 replicates, respectively, iterated maximally five times, using a grid size of 100, 110, and 120, and projections that maximized sample size against the number of segregating size (73 for mainland, 8 for Newfoundland), as outlined in Portik *et. al* (69). The most likely demographic model (lowest AIC) was then optimized to determine the best values of *N*_*uA*_, *N*_*u*1_, *N*_*u*2_, *τ*_1_, *s*, and *f* using a similar process outlined above except specifying explicit upper and lower bounds, using three-fold cross validation, and repeating the entire process five times, as detailed in Portik *et al.* (69). Model fit was assessed by comparing empirical model fit to 100 SFS simulations with two-fold cross validation (Fig. S6), as outlined in Barratt *et al.* (70). This process was again repeated to create a 1D SFS solely for Newfoundland to assess the presence of a historical bottleneck (Fig. S7). Split time was determined by first using the equation, (θ = 4*N*_*ref*_*µL*), which after solving for *N*_*ref*_ was used to determine divergence with the equation (*τ*_*i*_ = 2*τ*_1_*N*_*ref*_*G)*. To assess sensitivity (Table S4) we used 3 values of the neutral mutation rate (*µ* = 0.9 × 10^-9^, 1.1 × 10^-9^, and 1.3 × 10^-9^) as used in other related species (71), determined *L* from the total sequence assayed, used the value of *τ*1 directly from ∂a∂i, and assessed three different values of generation time in years (*G* = 1.8, 2.3, and 2.8) as observed in bobcat (*Lynx rufus*) generation times (72).

### Bioinformatics – DNA methylation

We identified methylated and non-methylated positions by first filtering BAM files for incomplete bisulfite conversion based on reads containing more than three methylated positions in a CHH or CHG context. In the remaining reads, methylated positions in a CpG context with a sequencing depth of at least five were extracted with Bismark (29), truncating the last two bases of the forward mate-paired reads (R1) and the first two bases of the reverse mate-paired (R2) reads. Methylation polymorphisms in areas of overlap between read pairs were extracted only once. After confirming that our non-directional library contained roughly equal reads for all possible amplified DNA strands, we proceeded to analysis.

We first generated a custom CpG island annotation track using hidden Markov models based on CpG_o/e_ implemented in *makeCGI* (73). Only islands with a calculated posterior probability greater than 99.5% were retained for analysis, based on CpG_o/e_. Mapped and extracted methylated sites were then imported into Seqmonk (74) using the generic text importer, and raw data were qualitatively visualized against the annotated domestic cat genome (felCat9.0). We analyzed DNA methylation over CpG islands and gene bodies by creating 5,000-bp running windows directly over and 25,000-bp upstream of gene bodies, combined with windows directly over CpG islands. Remaining, unannotated regions of the genome were then also divided into 5,000-bp windows, and added to the analysis. Each window was assigned a methylated percentage score based on the overall ratio of methylated to non-methylated bases within the feature. A window size of 5,000-bp was chosen based on maximizing a number of windows with all 95 individuals, and is similar to sizes seen in other studies (75). We filtered this window-set for regions that had at least one CpG and equal representation from each population due to low coverage of some individuals, although many windows contained more CpGs (Table S10). No overlap between SNPs and DMPs were identified (Fig. 2).

### Sensitivity analyses

To determine the implications of missing data and PCoA axis retention thresholds, we performed two sensitivity analyses. The first analysis examined the effects of missing data by repeating all analyses using a subset of the top 10 individuals per population (N = 40), and again with the top 6 individuals per population (*N* = 24), across both SNP and DNA methylation datasets (Fig. S1 – S5). Overall trends in explanatory effects (R^2^) and qualitative inferences (PCoA clustering) were investigated and no change in inferences were determined. We examined the implications of arbitrary PCoA axis retention by repeating all analyses, but instead using different cumulative variation explained thresholds as response variables. We performed a number of db-RDAs using all axes explaining 30%, 50%, 75%, and 95% cumulative variation as response variables, and results were qualitatively similar regardless of axis retention (Fig. S4). We also assessed if feature-specific patterns of DNA methylation were different between populations. We did this by treating our CpG island and gene body dataset separately from our unannotated region dataset, and repeating all analyses (Fig. S3 – S5). We also assessed where our DNA methylation dataset fell on the spectrum of missing at random (MAR) or missing not a random (MNAR) per locus between individuals (Fig. S11). Although a number of individuals in Newfoundland show low per locus sampling, this is likely due to stochastic inequal amplification or miscalibration of equimolar pooling during super-pooling of the final RRBS library, as roughly 12 sequential individuals (the number of individuals in a pool) suffered from low coverage, while remaining individuals show similar sampling to other populations (Fig. S11). We repeated ordinations after removal of these individuals, which showed no impact on observed patterns, suggesting that missing data is not the cause of the unique epigenetic structuring of Newfoundland (Fig. S11).

### Quantifying environmental associations

To determine if patterns of DNA methylation could be explained by macro-scale winter conditions, geographic distance, or insular divergence, we performed a distanced-based redundancy analysis on the summarized DNA methylation data. Meaningful axes explaining > 30% of the cumulative variation in the data were used as response variables in a distance-based redundancy analysis (db-RDA) conducted in *vegan* (76), using variables that putatively describe the environmental determinants of population structure in Canada lynx. Our covariates included a binary variable of insularity, which identified the Newfoundland population against mainland populations and was used to describe the largely impassable barrier of the Strait of Belle Isle between Newfoundland and mainland Labrador (77). A variable of geographic distance was included which was simply the first axis of a principal coordinates analysis (PCoA) on a Euclidean distance matrix of latitude and longitude (*PCo*1 = 99.7% of the variation, Fig. S12). In addition to the geographic variables, we included a biotic variable of percent tree cover (78), a randomly generated numerical variable to assess the effect of noise, and a climate variable describing winter conditions. To create the climate variable, we performed a PCA on climate data to prevent multi-collinearity, which reduced annual temperature ranges, winter precipitation, and minimum coldest temperature to a single PCA axis (28) (*PC*1 = 85.6% of the variation, Fig. S13). Linearity was confirmed between response and explanatory variables, and multi-collinearity between explanatory variables was assessed using the VIF and any variables > 4 were removed (Table S6). Step-wise model selection using the function *ordistep* within *vegan* (76) was performed to isolate the best overall model using a QR decomposition technique based on p-values (Table S5). To isolate the individual explanatory power of each variable, we performed partial distanced-based redundancy analyses (p-dbRDAs) on the variables that were identified as significant in the full db-RDA.

### Differentially methylated regions and gene ontology

We identified differentially methylated regions with functional biological correlates by performing beta-regressions on windows over CpG islands and gene bodies for all 95 individuals with percent methylation as the response variable and population as the explanatory variable. Beta regressions are appropriate for proportion or percentage data (79), and we set an alpha threshold at conservative levels seen in similar studies (80) (*p* < 0.001). Direct overlap between our differentially methylated regions and the felcat9.0 gene annotations were extracted using Seqmonk (74). Samples were then categorized as ‘Mainland’ or ‘Newfoundland’ and population averages and standard error of the mean were calculated for visualization. We then identified specific enrichment for biological functions of differentially methylated genes using the Panther-GO Gene Ontology Consortium database (http://geneontology.org/page/go-enrichment-analysis) with significance thresholds seen in other studies (80).

## ACKNOWLEDGEMENTS

The authors thank Niels C.A.M. Wagemaker for his assistance with library preparation and his generous donation of methylated adapters. We would also like to thank Felix Krueger and Sibelle Torres Vilaça for their assistance with bioinformatics. We would like to thank Joe M. Northrup (OMNR), Elizabeth M. Kirepka, Lynne E. Beaty, Angela R. Eads, and Melanie R. Boudreau for their invaluable statistical advice and friendly manuscript reviews. We thank the Natural Resources DNA Profiling and Forensic Centre (NRDPFC) for their assistance with extractions and laboratory equipment, and the Centre for Applied Genomics with sequencing troubleshooting. This work was funded by the CFI-JELF (358-47; A.B.A.S.), ComputeCanada (GME-665-01; A.B.A.S.), NSERC Discovery Grants (D.L.M., A.B.A.S.) and CREATE-enviro (J.B.J.).

## Author Contributions

J.B.J., D.L.M., and A.B.A.S. designed the study; J.B.J. performed research and analyzed data; J.B.J. wrote the manuscript with input from A.B.A.S. and D.L.M.

## Materials and Correspondence

Inquiries can be directed to the corresponding author, J.B.J.

## REFERENCES

1. Beaumont MA, Balding DJ (2004) Identifying adaptive genetic divergence among populations from genome scans. Mol Ecol 13(4):969–980.

2. Stinchcombe JR, Hoekstra HE (2008) Combining population genomics and quantitative genetics: finding the genes underlying ecologically important traits. Heredity (Edinb) 100(2):158–170.

3. Shafer ABA, Wolf JBW (2013) Widespread evidence for incipient ecological speciation: A meta-analysis of isolation-by-ecology. Ecol Lett 16(7):940–950.

4. Feder JL, Egan SP, Nosil P (2012) The genomics of speciation-with-gene-flow. Trends Genet 28(7):342–350.

5. Jones PA (2001) The Role of DNA Methylation in Mammalian Epigenetics. Science 293(5532):1068–1070.

6. Dubin MJ, et al. (2015) DNA methylation in Arabidopsis has a genetic basis and shows evidence of local adaptation. Elife 4(5):1–23.

7. Artemov A V., et al. (2017) Genome-Wide DNA Methylation Profiling Reveals Epigenetic Adaptation of Stickleback to Marine and Freshwater Conditions. Mol Biol Evol 34(9):2203–2213.

8. Lea AJ, Altmann J, Alberts SC, Tung J (2016) Resource base influences genome-wide DNA methylation levels in wild baboons (Papio cynocephalus). Mol Ecol 25(8):1681–1696.

9. Fujita N, et al. (2003) Methyl-CpG binding domain 1 (MBD1) interacts with the Suv39h1-HP1 heterochromatic complex for DNA methylation-based transcriptional repression. J Biol Chem 278(26):24132–24138.

10. Lorincz MC, Dickerson DR, Schmitt M, Groudine M (2004) Intragenic DNA methylation alters chromatin structure and elongation efficiency in mammalian cells. Nat Struct Mol Biol 11(11):1068–1075.

11. Maunakea AK, et al. (2010) Conserved role of intragenic DNA methylation in regulating alternative promoters. Nature 466(7303):253–257.

12. Wutz A, P. Barlow D (1998) Imprinting of the mouse Igf2r gene depends on an intronic CpG island. Mol Cell Endocrinol 140(1–2):9–14.

13. Han L, Su B, Li W-H, Zhao Z (2008) CpG island density and its correlations with genomic features in mammalian genomes. Genome Biol 9(5):R79.

14. Jones PA (2012) Functions of DNA methylation: Islands, start sites, gene bodies and beyond. Nat Rev Genet 13(7):484–492.

15. Row JR, et al. (2012) Dispersal promotes high gene flow among Canada lynx populations across mainland North America. Conserv Genet 13(5):1259–1268.

16. Prentice MB, et al. (2017) Selection and drift influence genetic differentiation of insular Canada lynx (Lynx canadensis) on Newfoundland and Cape Breton Island. Ecol Evol 7(9):3281–3294.

17. Stenseth NC, et al. (1999) Common Dynamic Structure of Canada Lynx Populations Within Three Climatic Regions. Science 285(5430):1071–1073.

18. Row JR, et al. (2014) The subtle role of climate change on population genetic structure in Canada lynx. Glob Chang Biol 20(7):2076–2086.

19. Van Zyll De Jong CG (1975) Differentiation of the Canada lynx, Felis (Lynx) canadensis subsolana, in Newfoundland. Can J Zool 53(6):699–705.

20. Khidas K, Duhaime J, Huynh HM (2013) Morphological Divergence of Continental and Island Populations of Canada Lynx. Northeast Nat 20(4):587–608.

21. Foster JB (1964) Evolution of mammals on Islands. Nature 202(4929):234–235.

22. Van Valen L (1973) Pattern and the balance of nature. Evol Theory 1(1):31–49.

23. Wayne RK, et al. (1991) A morphologic and genetic study of the Island fox, Urocyon littoralis. Evolution (N Y) 45(8):1849–1869.

24. Lomolino M V. (2005) Body size evolution in insular vertebrates: Generality of the island rule. J Biogeogr 32(10):1683–1699.

25. Flatscher R, Frajman B, Schönswetter P, Paun O (2012) Environmental Heterogeneity and Phenotypic Divergence: Can Heritable Epigenetic Variation Aid Speciation? Genet Res Int 2012:1–9.

26. Stenseth NC, et al. (2004) Snow conditions may create an invisible barrier for lynx. Proc Natl Acad Sci 101(29):10632–10634.

27. Keller TE, Lasky JR, Yi S V. (2016) The multivariate association between genomewide DNA methylation and climate across the range of Arabidopsis thaliana. Mol Ecol 25(8):1823–1837.

28. Fick SE, Hijmans RJ (2017) WorldClim 2: new 1-km spatial resolution climate surfaces for global land areas. Int J Climatol 37(12):4302–4315.

29. Krueger F, Andrews SR (2011) Bismark: A flexible aligner and methylation caller for Bisulfite-Seq applications. Bioinformatics 27(11):1571–1572.

30. Gutenkunst RN, Hernandez RD, Williamson SH, Bustamante CD (2009) Inferring the Joint Demographic History of Multiple Populations from Multidimensional SNP Frequency Data. PLoS Genet 5(10):e1000695.

31. South GR (1983) Biogeography and ecology of the Islands of Newfoundland (Dr. W. Junk Publishers, The Hague).

32. Rueness EK, et al. (2003) Ecological and genetic spatial structuring in the Canadian lynx. Nature 425:69–72.

33. Chatterjee TK, et al. (2014) HDAC9 knockout mice are protected from adipose tissue dysfunction and systemic metabolic disease during high-fat feeding. Diabetes 63(1):176–187.

34. Weber KL, Fischer RS, Fowler VM (2007) Tmod3 regulates polarized epithelial cell morphology. J Cell Sci 120(20):3625–3632.

35. Conley CA, Fritz-Six KL, Almenar-Queralt A, Fowler VM (2001) Leiomodins: Larger Members of the Tropomodulin (Tmod) Gene Family. Genomics 73(2):127–139.

36. Saykally JN, Dogan S, Cleary MP, Sanders MM (2009) The ZEB1 Transcription Factor Is a Novel Repressor of Adiposity in Female Mice. PLoS One 4(12):e8460.

37. Taudt A, Colomé-Tatché M, Johannes F (2016) Genetic sources of population epigenomic variation. Nat Rev Genet 17(6):319–332.

38. Vidalis A, et al. (2016) Methylome evolution in plants. Genome Biol 17(1):1–14.

39. Ichikawa K, et al. (2017) Centromere evolution and CpG methylation during vertebrate speciation. Nat Commun 8(1). doi:10.1038/s41467-017-01982-7.

40. Yu R, Wang X, Moazed D (2018) Epigenetic inheritance mediated by coupling of RNAi and histone H3K9 methylation. Nature 558(7711):615–619.

41. He L, et al. (2018) A naturally occurring epiallele associates with leaf senescence and local climate adaptation in Arabidopsis accessions. Nat Commun 9(1):1–11.

42. Vogt G (2017) Facilitation of environmental adaptation and evolution by epigenetic phenotype variation: insights from clonal, invasive, polyploid, and domesticated animals. Environ Epigenetics 3(1):1–17.

43. Vogt G (2018) Investigating the genetic and epigenetic basis of big biological questions with the parthenogenetic marbled crayfish: A review and perspectives. J Biosci 43(1):189–223.

44. Greenspoon PB, Spencer HG (2018) The evolution of epigenetically mediated adaptive transgenerational plasticity in a subdivided population. Evolution (N Y):1–8.

45. Joo JE, et al. (2018) Heritable DNA methylation marks associated with susceptibility to breast cancer. Nat Commun 9(1):867.

46. Liu G, Wang W, Hu S, Wang X, Zhang Y (2018) Inherited DNA methylation primes the establishment of accessible chromatin during genome activation. Genome Res:gr.228833.117.

47. Colome-Tatche M, et al. (2012) Features of the Arabidopsis recombination landscape resulting from the combined loss of sequence variation and DNA methylation. Proc Natl Acad Sci 109(40):16240–16245.

48. Kazachenka A, et al. (2018) Identification, Characterization, and Heritability of Murine Metastable Epialleles: Implications for Non-genetic Inheritance. Cell 175(5):1259–1271.e13.

49. Schmitz RJ, et al. (2013) Epigenome-wide inheritance of cytosine methylation variants in a recombinant inbred population. Genome Res 23(10):1663–1674.

50. Jeremias G, et al. (2018) Synthesizing the role of epigenetics in the response and adaptation of species to climate change in freshwater ecosystems. Mol Ecol 27(13):2790–2806.

51. Verhoeven KJF, VonHoldt BM, Sork VL (2016) Epigenetics in ecology and evolution: What we know and what we need to know. Mol Ecol 25(8):1631–1638.

52. van Gurp TP, et al. (2016) epiGBS: reference-free reduced representation bisulfite sequencing. Nat Methods 13(4):322–324.

53. Raine A, Liljedahl U, Nordlund J (2018) Data quality of whole genome bisulfite sequencing on Illumina platforms. PLoS One 13(4):e0195972.

54. Andrews S (2010) FastQC: A quality control tool for high throughput sequence data. Available at https://www.bioinformatics.babraham.ac.uk/projects/fastqc/. Online.

55. Martin M (2011) Cutadapt removes adapter sequences from high-throughput sequencing reads. EMBnet.journal 17(1):10–12.

56. Krueger F (2012) Trim Galore! : A wrapper tool around Cutadapt and FastQC to consistently apply quality and adapter trimming to FastQ files. Available at https://www.bioinformatics.babraham.ac.uk/projects/trim_galore/. Online.

57. Kobayashi H, et al. (2016) Repetitive DNA methylome analysis by small-scale and singlecell shotgun bisulfite sequencing. Genes to Cells 21(11):1209–1222.

58. Abascal F, et al. (2016) Extreme genomic erosion after recurrent demographic bottlenecks in the highly endangered Iberian lynx. Genome Biol 17(1):251.

59. Kunde-Ramamoorthy G, et al. (2014) Comparison and quantitative verification of mapping algorithms for whole-genome bisulfite sequencing. Nucleic Acids Res 42(6):e43–e43.

60. Guo W, et al. (2017) CGmapTools improves the precision of heterozygous SNV calls and supports allele-specific methylation detection and visualization in bisulfite-sequencing data. Bioinformatics 34(February):381–387.

61. Lea AJ, Vilgalys TP, Durst PAP, Tung J (2017) Maximizing ecological and evolutionary insight in bisulfite sequencing data sets. Nat Ecol Evol 1(8):1074–1083.

62. Danecek P, et al. (2011) The variant call format and VCFtools. Bioinformatics 27(15):2156–2158.

63. Maechler M, Rousseeuw P, Struyf A, Hubert M, Hornik K (2018) cluster: Cluster Analysis Basics and Extensions. R package version 2.0.7-1.

64. R Core Team (2017) R: A language and environment for statistical computing. R Foundation for Statistical Computing, Vienna, Austria. URL https://www.R-project.org/. Online.

65. Jombart T (2008) Adegenet: A R package for the multivariate analysis of genetic markers. Bioinformatics 24(11):1403–1405.

66. Foll M, Gaggiotti O (2008) A genome-scan method to identify selected loci appropriate for both dominant and codominant markers: A Bayesian perspective. Genetics 180(2):977–993.

67. Kamvar ZN, Tabima JF, Grünwald NJ (2014) Poppr : an R package for genetic analysis of populations with clonal, partially clonal, and/or sexual reproduction. PeerJ 2:e281.

68. Excoffier L, Smouse P, Quattro J (1992) Analysis of Molecular Variance Inferred From Metric Distances Among DNA Haplotypes: Application to Human Mitochondrial DNA Restriction Data Laurent. Genetics 131(2):479–491.

69. Portik DM, et al. (2017) Evaluating mechanisms of diversification in a Guineo-Congolian tropical forest frog using demographic model selection. Mol Ecol 26(19):5245–5263.

70. Barratt CD, et al. (2018) Vanishing refuge? Testing the forest refuge hypothesis in coastal East Africa using genome-wide sequence data for seven amphibians. Mol Ecol (February):4289–4308.

71. Cho YS, et al. (2013) The tiger genome and comparative analysis with lion and snow leopard genomes. Nat Commun 4(1):2433.

72. Reding D, Bronikowski A, Johnson W, Clark W (2012) Pleistocene and ecological effects on continental-scale genetic differentiation in the bobcat (Lynx rufus). Mol Ecol 21(12):3078–3093.

73. Wu H, Caffo B, Jaffee HA, Irizarry RA, Feinberg AP (2010) Redefining CpG islands using hidden Markov models. Biostatistics 11(3):499–514.

74. Andrews S (2007) Seqmonk: A tool to visualise and analyse high throughput mapped sequence data. Available at https://www.bioinformatics.babraham.ac.uk/projects/seqmonk/. Online.

75. Guo JU, et al. (2011) Neuronal activity modifies the DNA methylation landscape in the adult brain. Nat Neurosci 14(10):1345–1351.

76. Dixon P (2003) VEGAN, a package of R functions for community ecology. J Veg Sci 14(6):927–930.

77. Koen EL, Bowman J, Wilson PJ (2015) Isolation of peripheral populations of Canada lynx (Lynx canadensis). Can J Zool 93(7):521–530.

78. DeFries RS, Hansen MC, Townshend JRG, Janetos AC, Loveland TR (2000) A new global 1-km dataset of percentage tree cover derived from remote sensing. Glob Chang Biol 6(2):247–254.

79. Ferrari SLP, Cribari-Neto F (2004) Betaregression for modelling rates and proportions. J Appl Stat 31(7):799–815.

80. Le Luyer J, et al. (2017) Parallel epigenetic modifications induced by hatchery rearing in a Pacific salmon. Proc Natl Acad Sci 114(49):12964–12969.

