## Supplemental Information for "Genome-wide sampling suggests island divergence accompanied by cryptic epigenetic plasticity in Canada lynx"

*** Corresponding author**: J.B. Johnson

**This PDF file includes:**

Supplementary Information Text

Figs. S1 to S14

Tables S2 to S13

References for SI reference citations

**RRBS Library Preparation**

Epidermal tissue samples were collected from Canada lynx (*Lynx canadensis*) pelts from either North American Fur Auctions (Toronto, Ontario) or Fur Harvester's Auctions, Inc. (North Bay, Ontario). Pelts were only dried, and not subjected to chemical processing that may impact molecular analyses. Small subsamples were taken consistently from the left side of the back leg on each pelt and were stored in individually barcoded envelopes until DNA extraction. Approximately 20 mg of pelt was removed from this subsample and suspended in 200 µl of lysis buffer and 20 µl of proteinase K to break down proteins and cell walls in preparation for DNA extraction. DNA was then isolated using magnetic beads with a MagneSil Blood Genomic Max Yield System (Promega) on an automated Janus robot (Perkin-Elmer). DNA concentrations were quantified using a Quant-It Picogreen Assay and were subsequently standardized to 20 ng/µl.

Full protocol and additional details can be found in van Gurp *et al.*, (2016) (van Gurp et al. 2016). A total of 95 Canada lynx samples and 1 sample of non-methylated lambda phage DNA (Sigma-Aldrich: D3654) were digested overnight (17 hours) at 37°C in individual 40 µl reactions containing 4 µl NEB 3.1 buffer, 1 µl AseI (10 units/µl; NEB: R0526S), 2 µl NsiI (20 units/µl; NEB: R0127L), 12 µl of UltraPure distilled water (Invitrogen), and 20 µl of DNA (400 ng total). We added individually barcoded in-line methylated adapters (Supplemental Table S1) to each digest product in a reaction containing 6 µl T4 DNA Ligase buffer (10x), 2 µl T4 DNA ligase (2,000,000 units/mL; NEB: M0202M), 4 µl AseI adapter (600 pg/µl), 4 µl NsiI adapter (600 pg/µl), and 8 µl of UltraPure $H_2O$. This 60 µl reaction was placed in a thermal cycler for three hours at 22°C, and then left on ice overnight at 4°C, with no heat inactivation.

Individually-barcoded samples were then amalgamated into 8 pools with 12 samples in each pool using a QIAQuick PCR Purification Kit (Qiagen: 28106), with 10 µl of 3M sodium acetate (NaOAC) added to each pool to neutralize pH. Pools were eluted in 50 µl and were then size-selected using AMPure XP magnetic beads with a ratio of 0.8x and freshly prepared 80% ethanol (Beckman Coulter: A63881). Residual ethanol was dried by resting samples on a heat block at 56°C until a hairline fracture was seen in the beads. Pools were then eluted in 23 µl UltraPure $H_2O$. Nicks in the adapters were then repaired in a 25 µl reaction with 19.25 µl of size-selected library, 2.5 µ 5-mC-dNTP mix (10 mM; Zymo: D1030), 2.5 µl NEB buffer 3.0 (10x), and 0.75 µl DNA polymerase I (10 units/µl; NEB: M0209L). This reaction was carried out on a thermal cycler for 60 minutes at 15°C.

Pools were subjected to bisulfite conversion using an EZ DNA Methylation-Lightning Kit (Zymo: D5030T) in a reaction with 23 µl size-selected and repaired library and 149.5 µl conversion reagent in a thermal cycler with an initial step at 98°C for 8 minutes followed by 60 minutes at 54°C. The remaining steps were performed according to manufacturer's instructions with a 20-minute desulphonation time and a final elution in 12 µl. Bisulfite-converted DNA was amplified in triplicate PCRs with 2 µl of bisulfite-converted library, 5 µl KAPA HiFi Uracil+ (Roche: KK2802), 0.3 µl of a forward Illumina primer, 0.3 µl reverse Illumina primer (Table S11), and 2.4 µl UltraPure $H_2O$ for 20 cycles (Table S12).

Triplicate PCRs were combined using a QIAQuick PCR Purification Kit with 10 µl 3M NaOAC and a final elution in 50 µl. We quantified the concentration of pools with a Qubit 3.0 (ThermoFisher Scientific) and respective quantities were amalgamated into a superpool with equal contribution from each pool. The superpool was cleaned with AMPure XP magnetic beads with a 0.8x ratio and was eluted in 24 µl. Final library fragment distribution was quantified with on an Agilent Bioanalyzer 2100 (Fig. S9) and successful flowcell ligation was confirmed with qPCR. 125-bp paired-end sequencing was then performed on a single lane on Illumina HiSeq2500 at the Hospital for Sick Children (Toronto, Ontario, Canada).

### Genetic Population Structure

SNPs were identified using CGmapTools (Guo et al. 2017), and vcf files were converted using PGDSpider v2.1.1.3 (Lischer and Excoffier 2012) for use in BayeScan v2.0 (Foll and Gaggiotti 2008). We ran BayeScan on this SNP matrix with default settings (50,000 burn-in, 20 pilot runs with a length of 5,000 chains, and 5,000 outputted iterations) four times, with prior odds for the neutral model modified between runs to from 10, 100, and 1,000. Pair-wise $F_{ST}$ was calculated using VCFtools (Danecek et al. 2011) (Table S13). Outlier loci were assessed with BayeScan v2.0. Most likely historical demographic patterns were assessed with ∂a∂i (Gutenkunst et al. 2009). Parameter lower and upper bounds for optimization of the 2D joint SFS for $N_{uA}, N_{u1}, N_{u2}, \tau_1, s,$ and $f$ were 10, 5, 1, 10, 0.1, 0.25 and 2, 1, 0.05, 0.1, 0.01, 0.05, respectively. Empirical, optimized parameters for calculating divergence and used for simulations (Fig. S6) were 4.12, 4.82, 4.18, 0.39, 0.13, 0.09, and 0.1. Upper and lower bounds for optimization of the 1D SFS for Newfoundland (Fig. S7) were 15, 10, 5 and 0.01, 0.05, 0.1 for $N_{uB}, N_{uF}, \tau_1$, respectively.

### Methylation Calling and Bisulfite Conversion Efficiency

Methylated sites in a CpG context were called using the *Bismark* methylation extractor function after removing any reads that were likely candidates for incomplete bisulfite conversion (> 3 methylated sites in a CHH or CHG context) (Krueger and Andrews 2011).

This step was deemed necessary as our non-methylated lambda phage DNA control identified minor incomplete bisulfite conversion. These results reinforce our decision to pool samples prior to bisulfite conversion, so that any reaction inconsistencies will apply to all samples and universalize any biases. Furthermore, we strongly recommend that all bisulfite sequencing experiments employ the use of a non-methylated lambda phage DNA as we demonstrate that incomplete bisulfite conversion does occur despite rigorous adherence to manufacturer's protocols.

We assessed relative levels of methylation directly around all transcripts using *Seqmonk*. We discovered a global reduction in methylation < 5,000-bp upstream of transcripts, with average levels resuming outside of the putative promoter region, consistent with results around the TSS in other research (Fig. S14) (Laine et al. 2016).

**Sensitivity Analyses**

*Missing Data*. First, we assessed the impacts of missing data by calculating Euclidean dissimilarity matrices, performing principal coordinates analyses (PCoAs) for both datasets at varied levels of missingness. Filtering was done by removing individuals with the most amount of missing data, while keeping equal representation for all populations. We evaluated the impacts of completeness at three sample levels ($N = 95; N = 40; N = 24$). These data were summarized with distance matrices using the function *daisy* (Maechler et al. 2018) and visualized with a PCoA using the function *dudi.pco* (Jombart 2008) (Fig. S1).

We then exported the axes of these PCoAs that explained > 30% of the total variation and performed db-RDAs for the methylation data on these axes as a response variable to determine if this variation in missing data would modify our inferences. We found a linear relationship between the amount of the missing data and the total explanatory power of the model ($R^2$), with coefficients increasing as the level of missing data decreases (Fig. S2).

*Temporal variation.* We assessed temporal variation in methylation patterns by analyzing a subset of the data where one population was sampled across two years. Due to sample access, our Canada lynx samples were collected from 2008-2012 (Table S1). We performed a db-RDA using the axes summarizing methylation patterns from the Alaskan population ($n = 23$), which has individuals from both 2009 and 2010, with year as an independent explanatory variable. db-RDAs for DNA methylation data detected no significant or explanatory effect of year on describing methylation patterns (Fig. S3).

*Axis retention for db-RDAs*. We examined the ramifications of our threshold for axis retention for db-RDA response variables by performing db-RDAs with variable amounts of PCoA axes. We examined the effect on $R^2$ when using axes that explained 30%, 50%, 75%, and 95% of cumulative variation as response variables for the DNA methylation dataset. We found a linear relationship between $R^2$ and cumulative variation explained (Fig. S4), consistent with the notion that including axes which explain less individual variation will decrease overall model fit, likely owing to explaining noise in the data.

*Feature-based DNA methylation.* To determine if feature-specific patterns in our DNA methylation dataset might bias our results, we repeated all analyses indicated in the full manuscript on two subsets of DNA methylation. One subset of analyses was done on methylation over CpG islands and gene bodies, while the other was done over unannotated regions of the genome. Correlations between datasets was high ($\rho = 0.88$), and PCoAs and db-RDAs identified largely similar trends (Fig. S3 – S5).

**Environmental determinants of DNA methylation**

We assessed the relationship between biogeographical variables and our molecular data by performing db-RDAs with PCoA axes that explain > 30% of the cumulative variation as response variables. Our explanatory variables included a binary variable with *Newfoundland* and *Mainland* as predictors, to represent the likely impermeable aquatic barrier between the island of Newfoundland and the mainland. Our second variable was geographic distance, which served as a null model as we would expect distribution in allele frequencies to correlate with distance across the landscape. To summarize this variable in a single vector and eliminate collinearity between latitude and longitude, we performed a PCoA on latitude and longitude (Fig. S12). The first axis of the PCoA explained overall variation exceptionally ($PCo1 = 99.73\%$ of the variation). To summarize as much variation as possible in winter climatic patterns into a single variable, we performed a PCA on bioclimatic variables using *sdmtoolbox* (Fig. S13) (Brown et al. 2017). We chose three variables with suspected biological significance for Canada lynx (*L. canadensis)*, particularly emphasizing variables that correlate with winter conditions. We summarized three bioclimatic variables from WorldClim (Fick and Hijmans 2017) including the annual temperature range (BIO7), minimum temperature of the coldest month (BIO6), and precipitation of the coldest quarter (BIO19). The first axis of this PCA described a majority of the variation ($PC1 = 85.62\%$ of the variation). We also included a continuous variable of tree cover (DeFries et al. 2000), and a randomly-generated vector to use as a null variable to mimic noise. Environmental data was extracted from each georeferenced lynx sample using R v3.4.2 (R Core Team 2017).

We then performed step-wise model selection using the PCoA axes of molecular data as response variables against these biogeographical explanatory variables. Step-wise selection was performed in *vegan* (Dixon 2003) using the function *ordistep*, performing forward and reverse step selection with 100 steps and 10,000 permutations. Variables were included when their p-value fell below a specified threshold at 0.05 and were removed from the model when its p-value rose above 0.2 (Table S5). Collinearity between explanatory variables was assessed by calculating the variance inflation factor (VIF; Table S6).

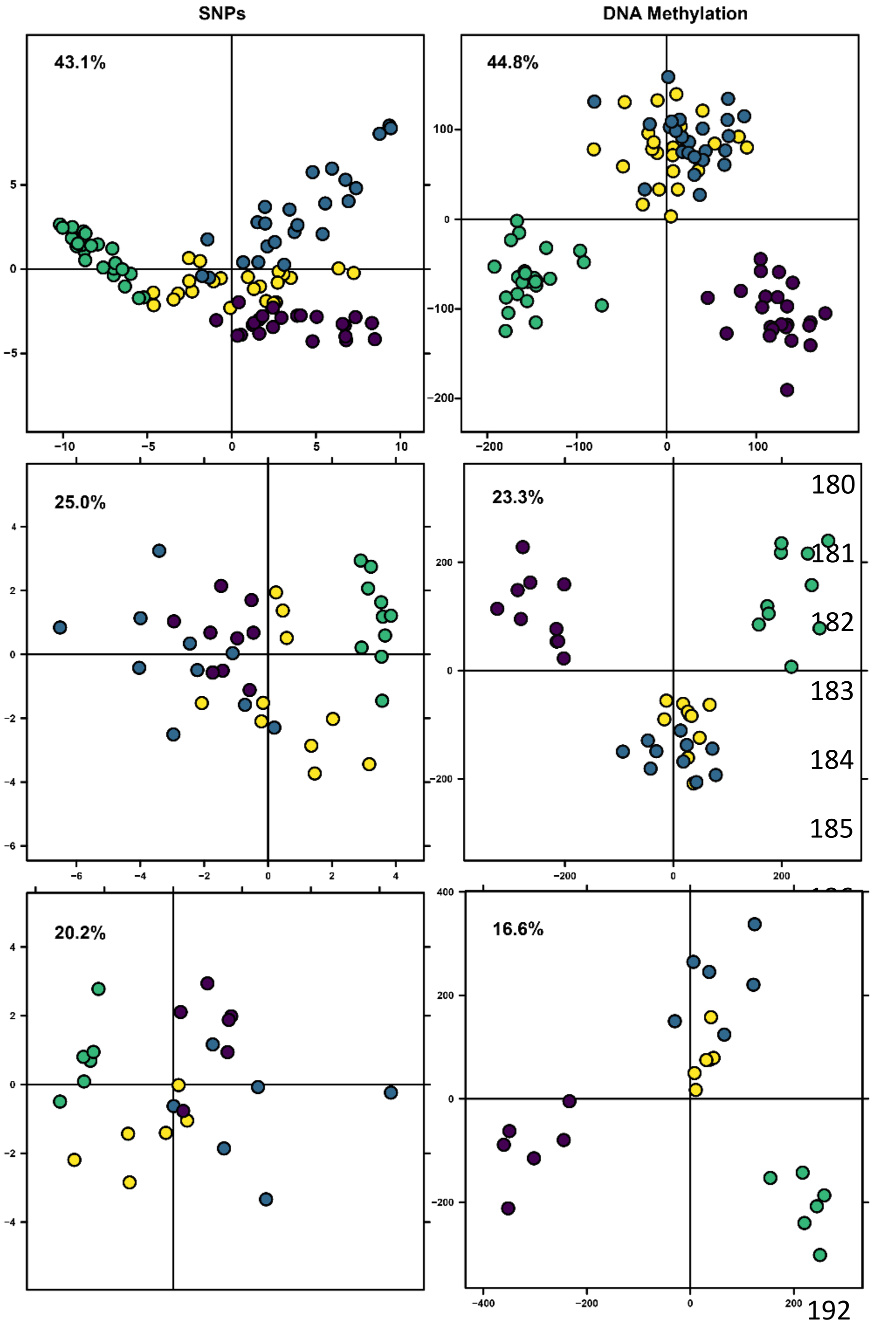


**Fig. S1.** Effects of missing data on observed patterns for SNP data (left) and DNA methylation data (right).

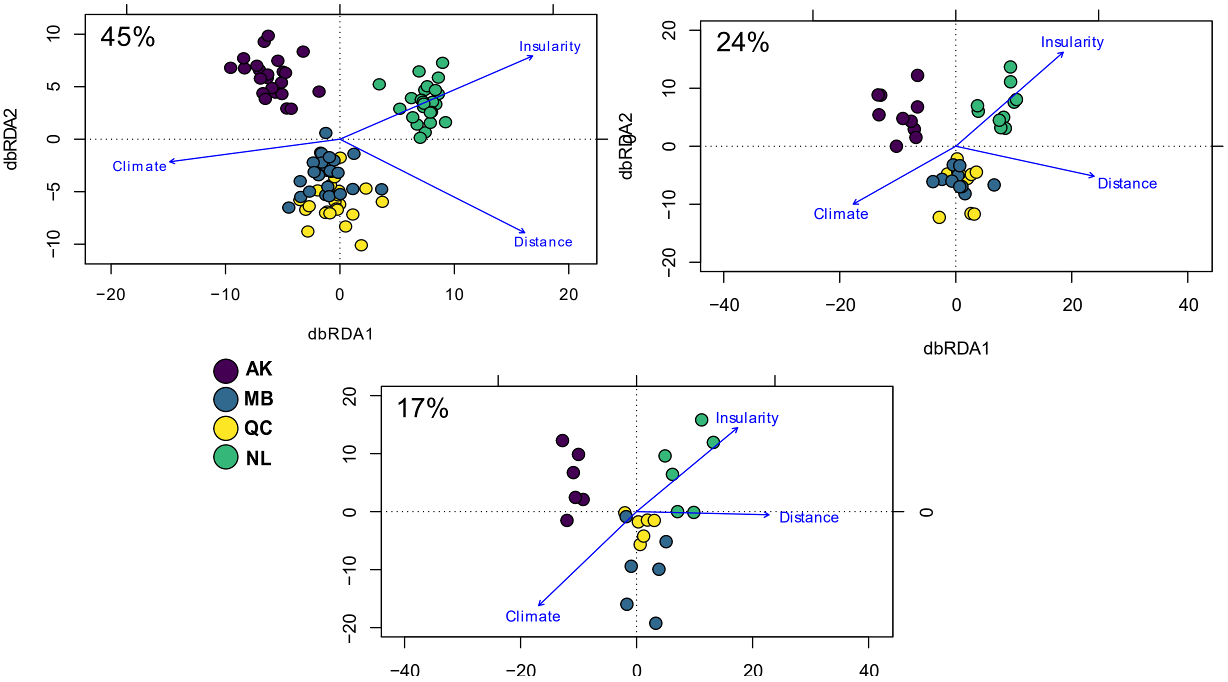


**Fig. S2.** Effects of missing data on db-RDA results for DNA methylation data, where percentage indicates the amount of missing data.

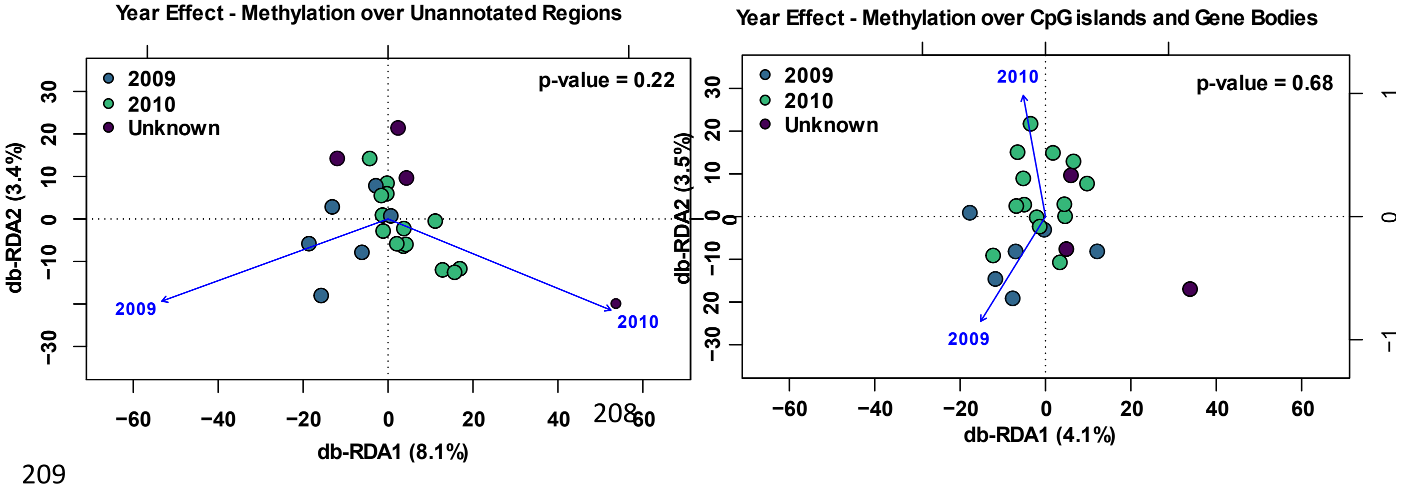


**Fig. S3.** Year effect on DNA methylation data for the Alaska population, where two different years of samples were available. This sensitivity analysis was divided into the two distinct subdivisions based on feature-type as described above in Supporting Information.

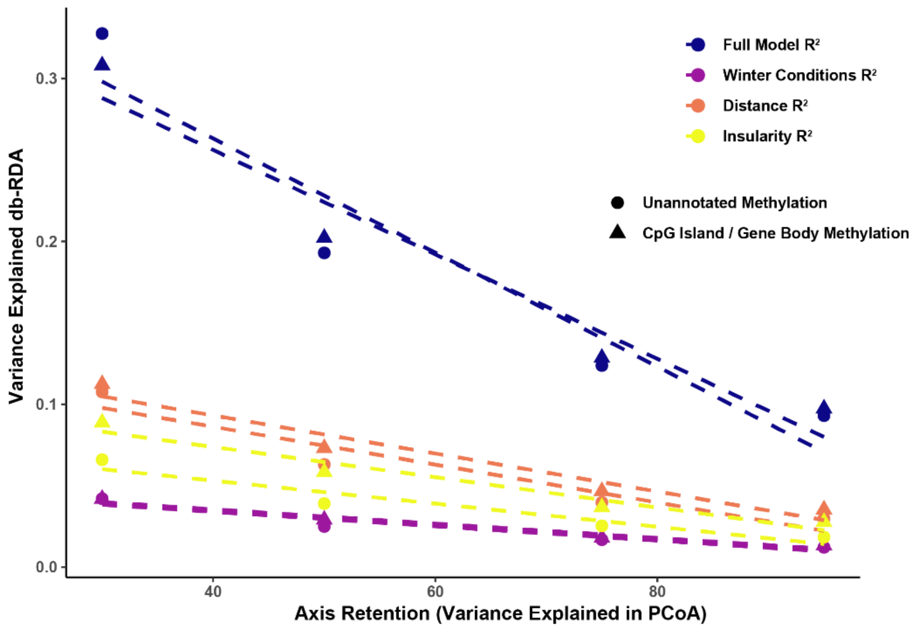


**Fig. S4.** Effect of axis retention on db-RDA results, where the X axis indicates the variation explained by the number of axes retained as explanatory variables for the db-RDA, and the Y axis indicates the $R^2$ explained by those axes. Biogeographical variables are indicated by color, where shapes describe the two DNA methylation datasets that were divided to assess feature-specific patterns.

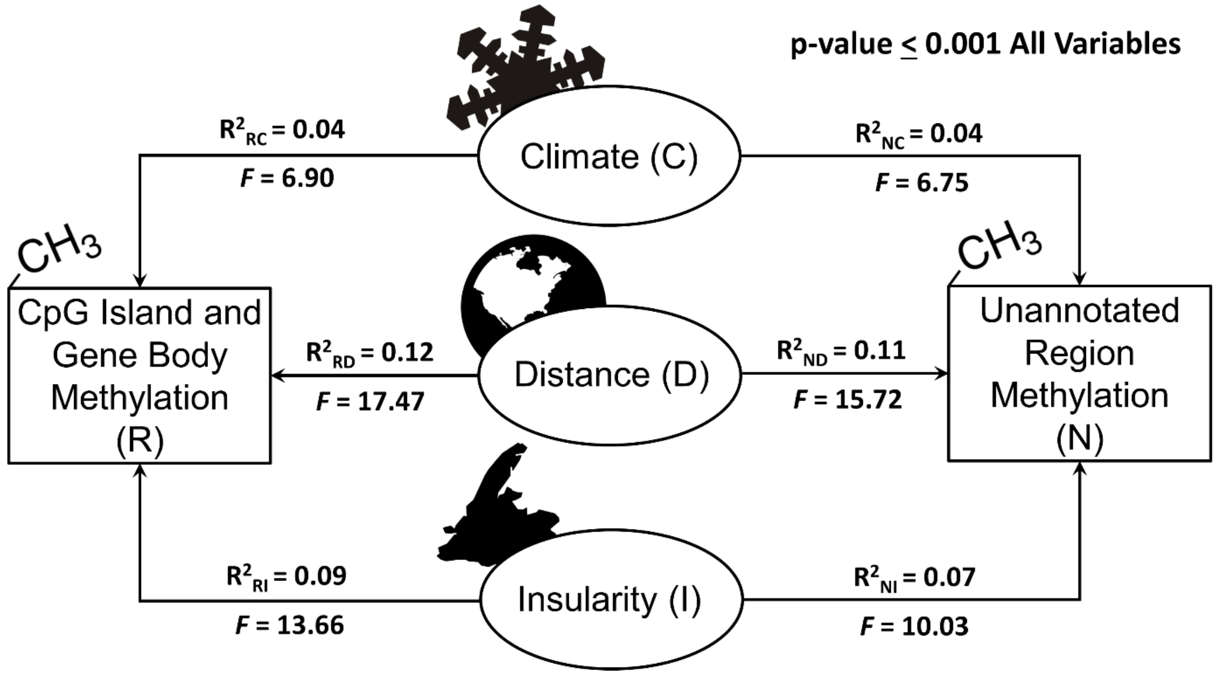


**Fig. S5.** Sensitivity analysis of p-db-RDA results between the feature-based subdivisions of DNA methylation data. The effect sizes indicate the independent explanatory effects of each variable on explaining methylation patterns, subtracted from the effect of any other variable. The effect size $R^2$ and the test-statistic is a pseudo-F generated using QR decomposition within vegan.

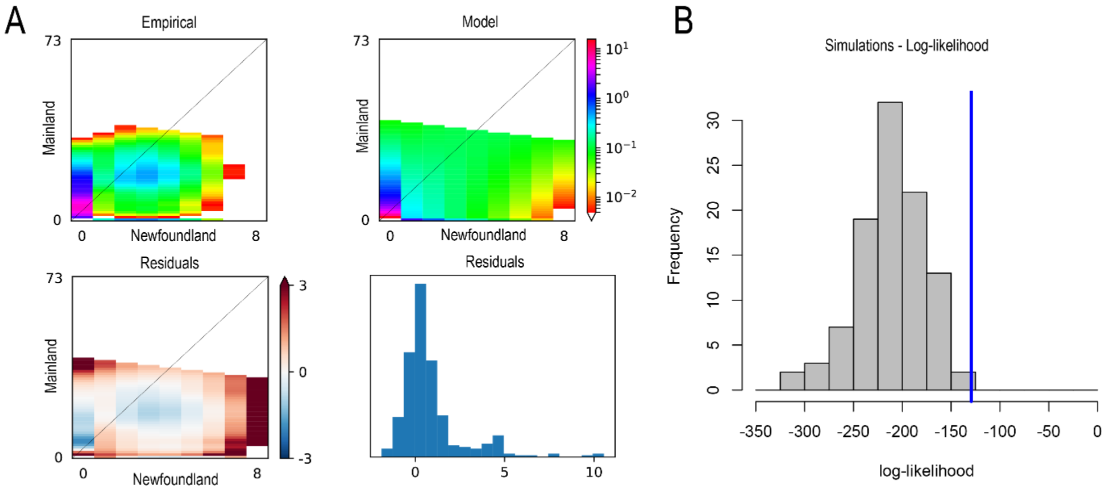


**Fig. S6.** Demographic inferences for Canada lynx using the site-frequency spectrum (SFS). **A.** Empirical (left) and model (right) joint-SFS for Canada lynx, created using ∂a∂i (Gutenkunst et al. 2009) with the model identified in Table S2. **B.** Assessing goodness of fit of our SFS against 100 simulations, using the procedure outlined in (Barratt et al. 2018).

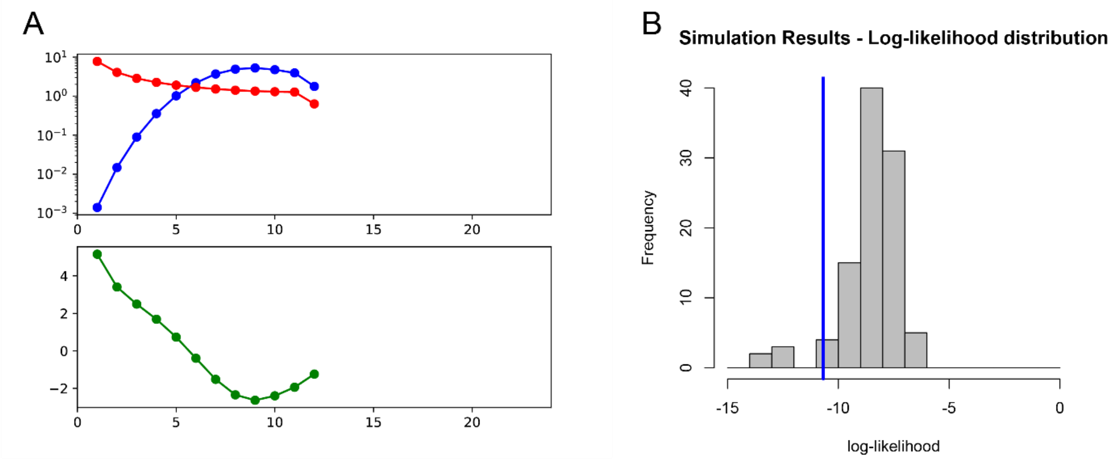


**Fig. S7.** Demographic inferences for Newfoundland Canada lynx using a 1-dimensional site-frequency spectrum (SFS). **A.** Model (red) and empirical data (blue), with residuals (green), created with a 1D-SFS *bottlegrowth* model using ∂a∂i (Gutenkunst et al. 2009) with the model identified in Table S2. **B.** Assessing goodness of fit of our 1-D SFS against 100 simulations, using the procedure outlined in (Barratt et al. 2018).

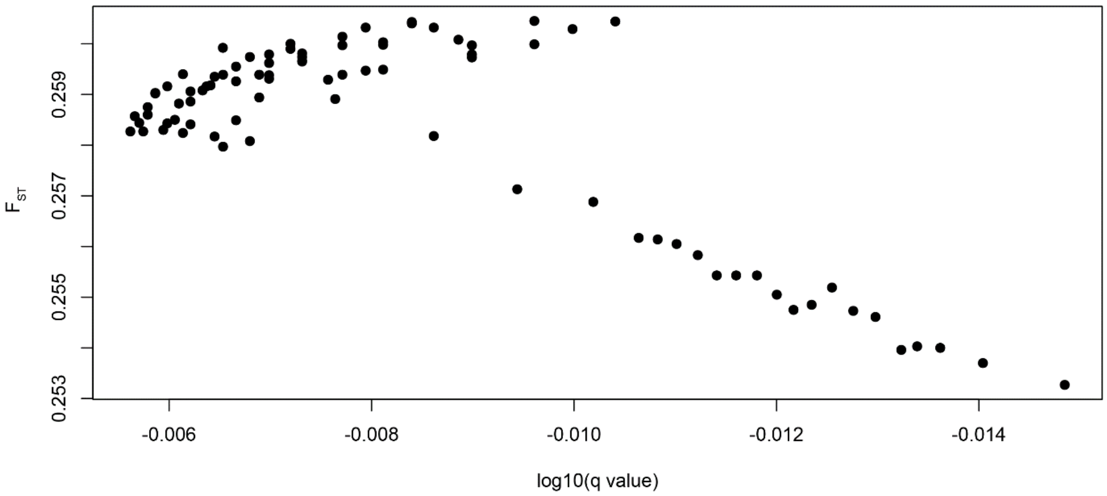


**Fig. S8.** Examining $F_{ST}$ outliers from SNP data to assess potential selection and confirm putatively neutral sites for characterizing population structure. Three runs were conducted to assess impact of prior odds (10, 100, 1000); above depicts a run with prior odds of 100.

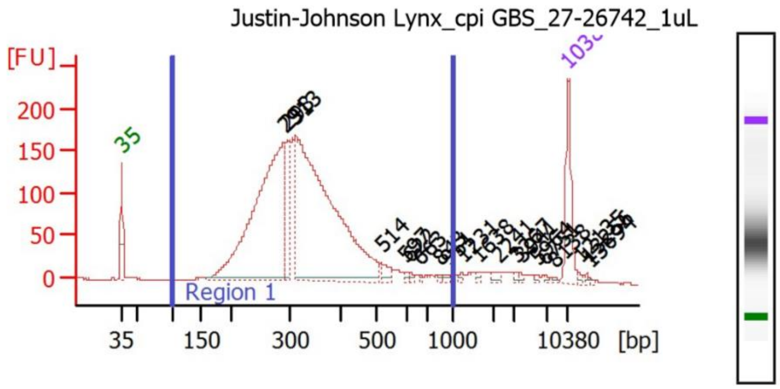


**Fig. S9.** Final library quality check for sequencing. Agilent Bioanalyzer 2100 results, showing final library fragment distribution before sequencing on a HiSeq 2500.

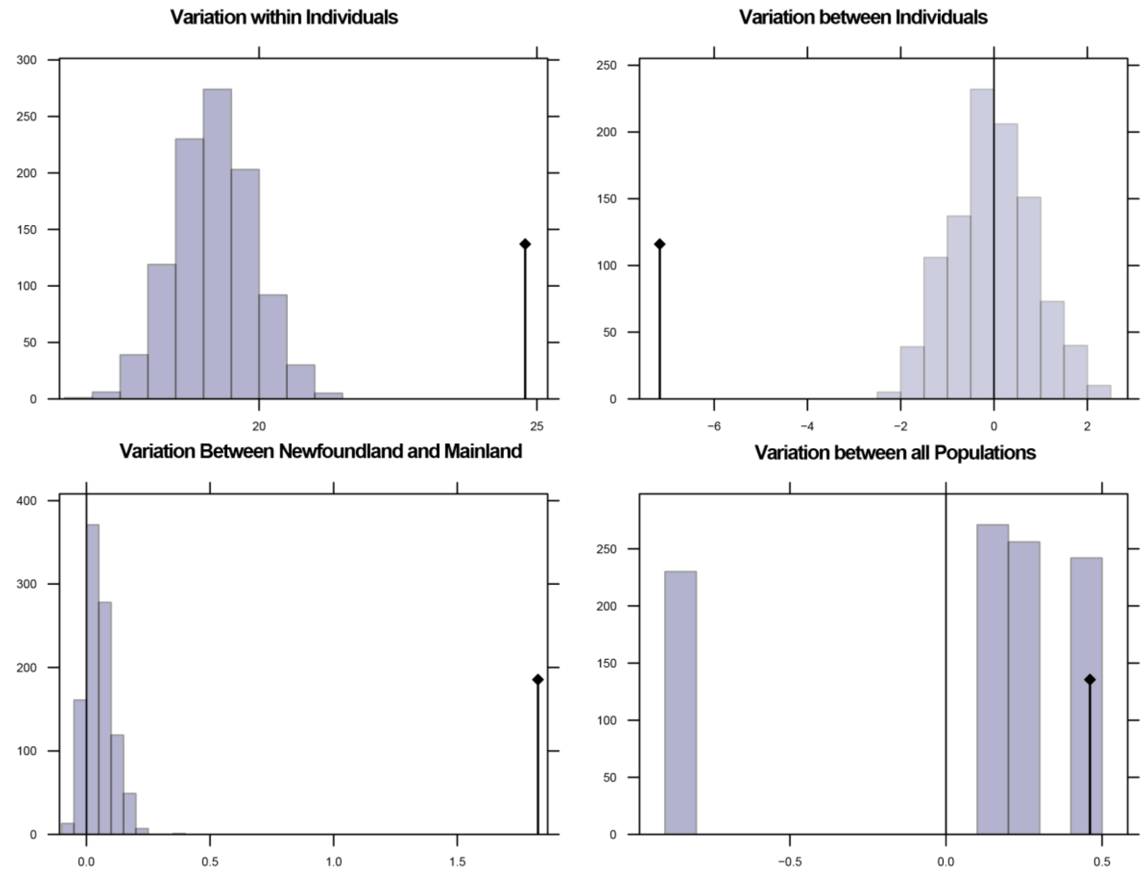


**Fig. S10.** Assessing population differentiation over putatively neutral SNPs (n = 85). Results of an AMOVA showing differentiation within individuals, between individuals, between Newfoundland and the mainland, and across all populations. No significant differentiation was identified across all populations, although differentiation was identified between Newfoundland and mainland.

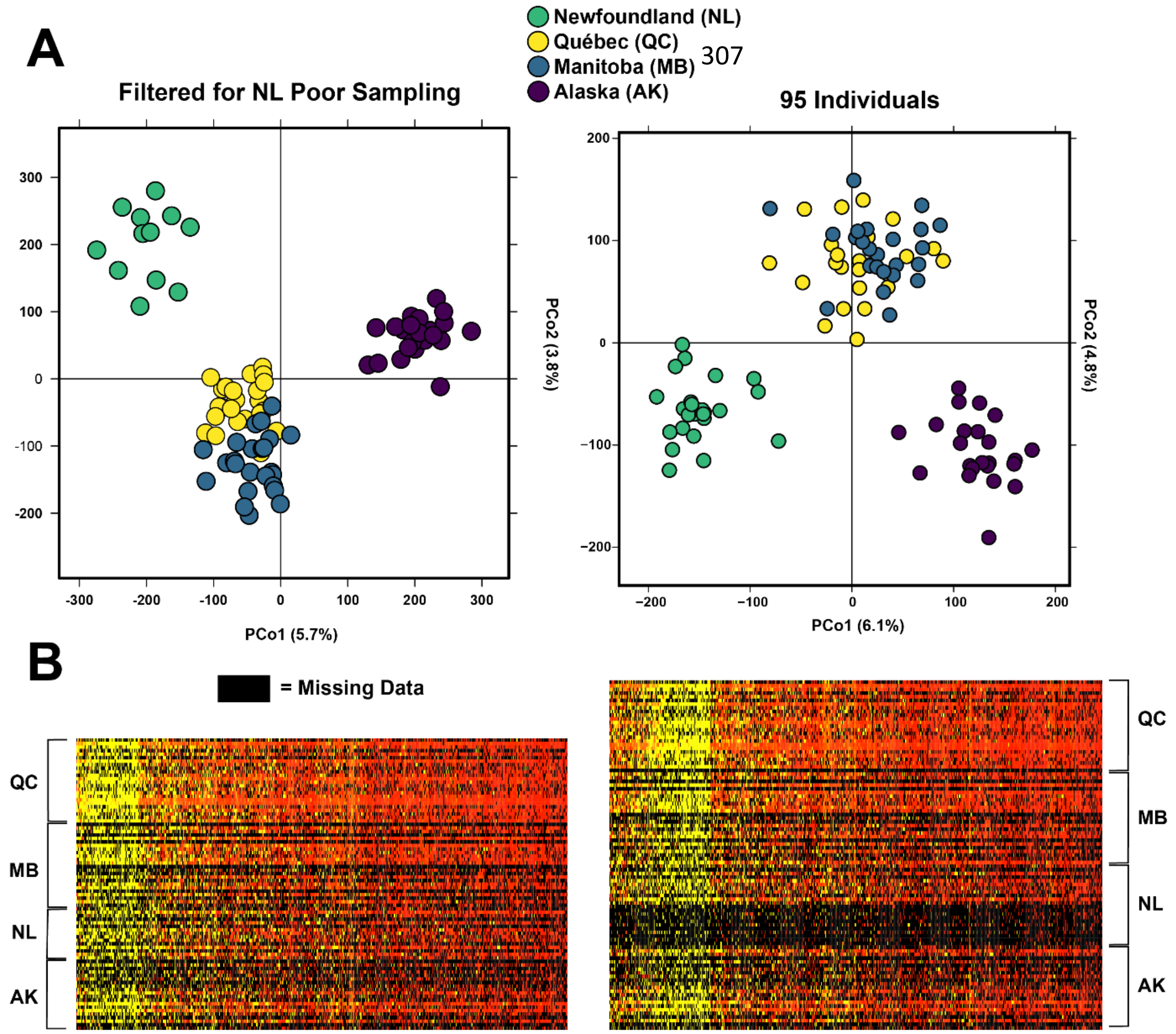


**Fig. S11.** Determining if observed patterns in DNA methylation between populations might be due to missing data that is either missing at random (MAR) or missing not at random (MNAR). **A)** PCoA filtered for 12 individuals from NL pool 6, which exhibited poor amplification (left), and full PCoA (right). **B)** Heatmap indicating missing data, with the 12 individuals from pool 6 exhibiting poor amplification (left), and full dataset (right).

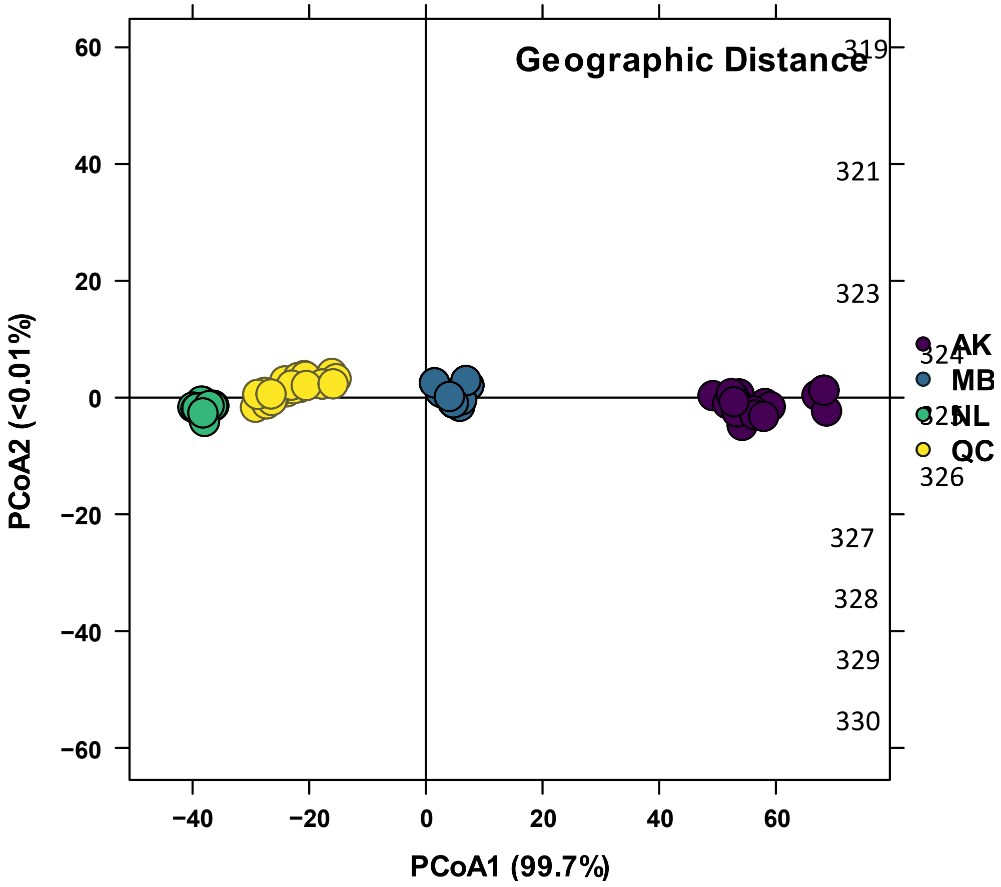


**Fig. S12.** PCoA summarizing a Euclidean distance matrix on geographic distance (latitude and longitude), used as a variable for db-RDAs to assess biogeographical relationships.

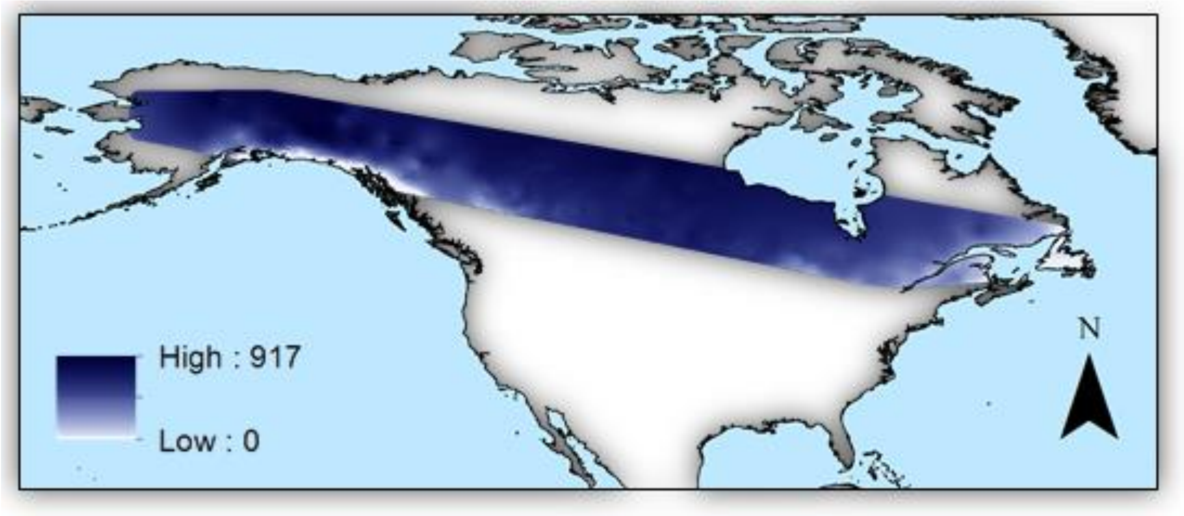


**Fig. S13.** PCA summarizing variation in winter climate across the study area. Variables included annual temperature range, minimum temperature of the coldest month, and precipitation of the coldest quarter, retrieved from WorldClim (Fick and Hijmans 2017). The first axis of this PCA explained 85.62% of the variation and was created to summarize winter conditions in a single vector for db-RDAs against molecular data.

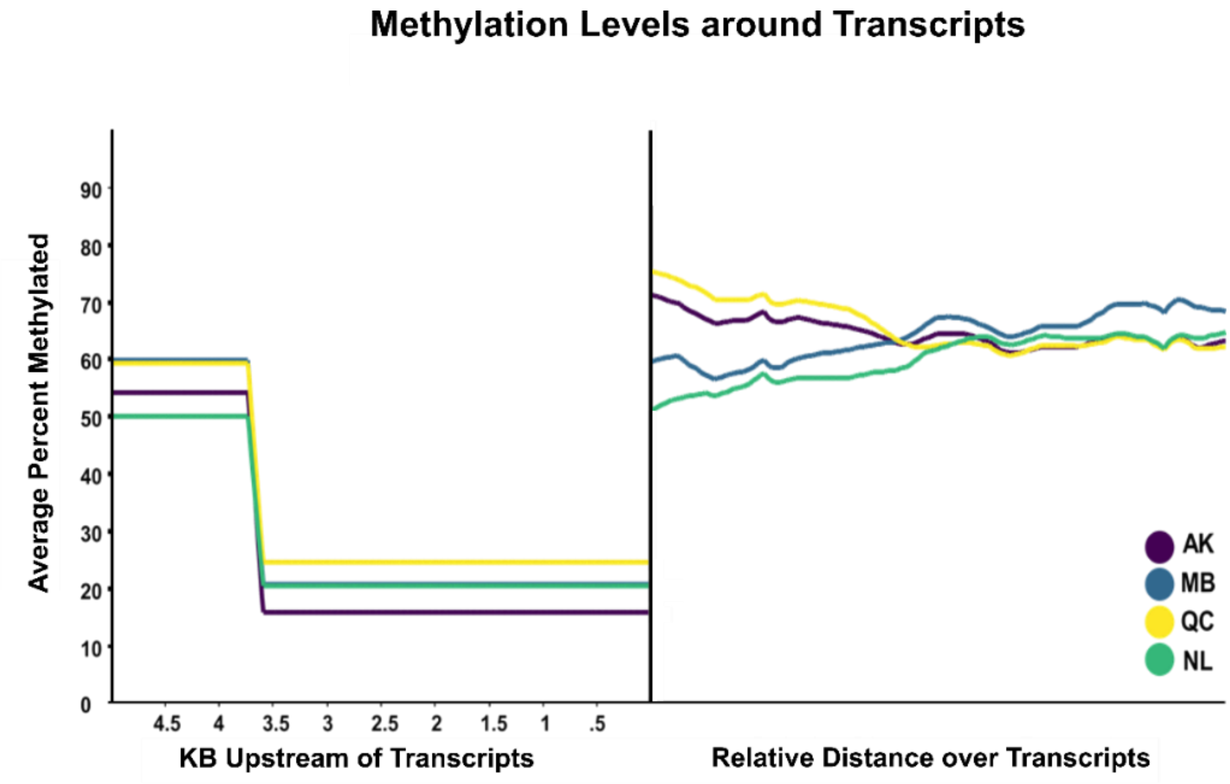


**Fig. S14.** Methylation levels around transcripts. The left pane shows methylation levels (%) 5,000-bp upstream of transcripts, while the right pane shows average methylation levels across the entire transcript

**Table S1.** (External File) Metadata for the Canada lynx samples used in this study.

**Table S2.** Total CpG positions sampled in our multiplexed RRBS library per individual.

| ID | Population | CpGs (Total) | CpGs (Methylated) | CpGs (Unmethylated) | ID | Population | CpGs (Total) | CpGs (Methylated) | CpGs (Unmethylated) |
|---|---|---|---|---|---|---|---|---|---|
| 1 | QC | 1559127 | 975249 | 583878 | 115 | NL | 848349 | 528533 | 319816 |
| 3 | QC | 9677599 | 5979220 | 3698379 | 116 | NL | 4584196 | 2823657 | 1760539 |
| 4 | QC | 3113980 | 1958791 | 1155189 | 117 | NL | 2218100 | 1342959 | 875141 |
| 7 | QC | 1086857 | 692770 | 394087 | 118 | NL | 574693 | 349525 | 225168 |
| 8 | QC | 4543922 | 2938438 | 1605484 | 119 | NL | 2380969 | 1444168 | 936801 |
| 10 | QC | 1133641 | 726212 | 407429 | 120 | NL | 743178 | 464795 | 278383 |
| 11 | QC | 4948967 | 3103424 | 1845543 | 122 | NL | 1869335 | 1121916 | 747419 |
| 14 | QC | 2252693 | 1410232 | 842461 | 124 | NL | 2081899 | 1263132 | 818767 |
| 15 | QC | 2351872 | 1506024 | 845848 | 125 | NL | 1709110 | 1019186 | 689924 |
| 16 | QC | 2310415 | 1478940 | 831475 | 126 | NL | 2210277 | 1321266 | 889011 |
| 18 | QC | 2620412 | 1674210 | 946202 | 127 | NL | 825325 | 504450 | 320875 |
| 24 | QC | 5645879 | 3482072 | 2163807 | 128 | NL | 3628796 | 2174464 | 1454332 |
| 26 | QC | 2263844 | 1363471 | 900373 | 130 | NL | 3557837 | 2250768 | 1307069 |
| 28 | QC | 1352291 | 827157 | 525134 | 131 | NL | 792658 | 501042 | 291616 |
| 29 | QC | 1304531 | 789471 | 515060 | 132 | NL | 497592 | 324460 | 173132 |
| 30 | QC | 1908462 | 1166563 | 741899 | 133 | NL | 1299273 | 855627 | 443646 |
| 32 | QC | 2348305 | 1450246 | 898059 | 134 | NL | 826448 | 539624 | 286824 |
| 33 | QC | 9325070 | 5701758 | 3623312 | 135 | NL | 466826 | 286417 | 180409 |
| 34 | QC | 7354904 | 4558329 | 2796575 | 136 | NL | 2145354 | 1414180 | 731174 |
| 35 | QC | 1327070 | 819659 | 507411 | 138 | NL | 387187 | 266953 | 120234 |
| 41 | QC | 3385709 | 2077010 | 1308699 | 139 | NL | 1360489 | 855721 | 504768 |
| 42 | QC | 1284472 | 785791 | 498681 | 141 | NL | 384821 | 260964 | 123857 |
| 43 | QC | 1448004 | 895200 | 552804 | 142 | NL | 768622 | 499995 | 268627 |
| 46 | QC | 2484592 | 1499599 | 984993 | 143 | NL | 789765 | 537244 | 252521 |

| | | | | | | | | | |
|---|---|---|---|---|---|---|---|---|---|
| 49 | MB | 262148 | 166675 | 95473 | 153 | AK | 1634938 | 1032900 | 602038 |
| 50 | MB | 2395053 | 1463126 | 931927 | 156 | AK | 7356939 | 4554355 | 2802584 |
| 51 | MB | 1751546 | 1070725 | 680821 | 160 | AK | 4148870 | 2518244 | 1630626 |
| 53 | MB | 358260 | 227506 | 130754 | 163 | AK | 634017 | 388458 | 245559 |
| 54 | MB | 3723357 | 2258657 | 1464700 | 166 | AK | 3898804 | 2340617 | 1558187 |
| 55 | MB | 399065 | 247231 | 151834 | 167 | AK | 1574355 | 1032507 | 541848 |
| 56 | MB | 4304444 | 2634259 | 1670185 | 170 | AK | 2403361 | 1487356 | 916005 |
| 58 | MB | 2392518 | 1469812 | 922706 | 171 | AK | 2193119 | 1351086 | 842033 |
| 61 | MB | 2094055 | 1256298 | 837757 | 172 | AK | 3711386 | 2325545 | 1385841 |
| 62 | MB | 2027636 | 1262476 | 765160 | 173 | AK | 1310251 | 817927 | 492324 |
| 64 | MB | 3446244 | 2092470 | 1353774 | 174 | AK | 2340507 | 1420282 | 920225 |
| 66 | MB | 4857321 | 3036124 | 1821197 | 175 | AK | 3385108 | 2069510 | 1315598 |
| 67 | MB | 671521 | 406397 | 265124 | 176 | AK | 3374440 | 2077908 | 1296532 |
| 69 | MB | 1779145 | 1067826 | 711319 | 177 | AK | 1989855 | 1209881 | 779974 |
| 71 | MB | 1311865 | 811108 | 500757 | 178 | AK | 1553655 | 940995 | 612660 |
| 73 | MB | 4698424 | 2839131 | 1859293 | 179 | AK | 6579509 | 3932904 | 2646605 |
| 74 | MB | 1806558 | 1121887 | 684671 | 180 | AK | 2190948 | 1352390 | 838558 |
| 77 | MB | 4794506 | 2890872 | 1903634 | 181 | AK | 7020902 | 4321079 | 2699823 |
| 79 | MB | 4856208 | 2988899 | 1867309 | 182 | AK | 2576543 | 1580942 | 995601 |
| 82 | MB | 703005 | 434135 | 268870 | 184 | AK | 1022789 | 613823 | 408966 |
| 83 | MB | 4725507 | 2886191 | 1839316 | 185 | AK | 3441754 | 2125182 | 1316572 |
| 85 | MB | 1145749 | 710171 | 435578 | 186 | AK | 511733 | 314526 | 197207 |
| 87 | MB | 3721534 | 2283752 | 1437782 | 187 | AK | 1405213 | 870422 | 534791 |
| 92 | MB | 1840085 | 1139925 | 700160 | | | | | |

**Table S3.** Results of the demographic model selection, determined using a joint-SFS with ∂a∂i (Gutenkunst et al. 2009) using a procedure detailed in Portik *et al.* (Portik et al. 2017).

| Model | log-likelihood | AIC | $X^2$ | Θ |
|---|---|---|---|---|
| Founder: No Migration, Early Admixture | -125.85 | 263.7 | 61.33 | 3.1 |
| Founder: Asymmetric Migration | -126.8 | 267.6 | 65.25 | 3.32 |
| Founder: Symmetric Migration | -128.99 | 269.98 | 68.31 | 3.25 |
| Founder: No Migration | -130.95 | 271.9 | 72.72 | 2.87 |
| Founder: No Migration, Late Admixture | -131.44 | 274.88 | 74.09 | 5.05 |
| Vicariance: Secondary Contact, Asymmetric Migration | -132.31 | 280.62 | 78.54 | 7.28 |
| Vicariance: Two Epoch Admixture Event | -133.85 | 281.7 | 82.19 | 7.26 |
| Vicariance: No Migration | -137.94 | 285.88 | 90 | 14.11 |
| Vicariance: Ancestral Asymmetric Migration | -135.31 | 286.62 | 85.69 | 4.17 |
| Vicariance: No Migration Late Admixture | -138.21 | 288.42 | 93.38 | 17.7 |
| Founder: No Migration Two Epoch Admixture Event | -138.53 | 291.06 | 90.63 | 9.59 |
| Vicariance: No Migration Early Admixture | -141.14 | 294.28 | 105.06 | 14.45 |

**Table S4.** Optimized demographic parameters for a 2D SFS indicating a founder effect with early asymmetric admixture into Newfoundland. $N_{uA}$ = Ancient Ne, $N_{u1}$ = Mainland Ne, $N_{u2}$ = Newfoundland Ne, $\tau_1$ = divergence time, $s$ = fraction of $N_{uA}$ to Newfoundland, $f$ = fraction of Newfoundland from $N_{u1}$, $L$ = sequence assayed, $\mu$ = neutral substitute rate, $G$ = generation time in years. Confidence intervals for $\tau_i$ indicated in matrix.

| ϴ | $N_{uA}$ | $N_{u1}$ | $N_{u2}$ | $\tau_1$ | $s$ | $f$ | $L$ |
|---|---|---|---|---|---|---|---|
| 4.12 | 4.82 | 4.18 | 0.39 | 0.13 | 0.09 | 0.1 | 21250 |

| | | | $G$ | |
|---|---|---|---|---|
| | | 1.8 | 2.3 | 2.8 |
| $\mu$ | 9E-10 | 24892.8 | 31807.5 | 38722.2 |
| | 1.10E-09 | 20366.9 | 26024.3 | 31681.8 |
| | 1.30E-09 | 17233.5 | 22020.6 | 26807.7 |

**Table S5.** db-RDA and p-db-RDA results for DNA methylation data, conducted in *vegan* (Dixon 2003). Axes of a PCoA describing DNA methylation data was used a response variable to determine environmental associations. Three variables were identified as significant (distance, insularity, winter climate), which were analyzed further in p-db-RDAs. Tree cover and a null RNG variable were not significant.

| Model | $F$ | ANOVA | $R^2$ | RDA1% | RDA2% | RDA1 ($F$) | RDA2 ($F$) | Insular | Distance | Climate | Tree Cover | Null |
|---|---|---|---|---|---|---|---|---|---|---|---|---|
| p-db-RDA: Distance | 17.07 | 0.001 | 0.121 | - | - | - | - | - | - | - | - | - |
| p-db-RDA: Insularity | 13.53 | 0.001 | 0.096 | - | - | - | - | - | - | - | - | - |
| p-db-RDA: Climate | 7.11 | 0.001 | 0.051 | - | - | - | - | - | - | - | - | - |
| Step-Wise Selection | - | - | - | - | - | - | - | 0.001 | 0.001 | 0.001 | 0.7685 | 0.7061 |
| Final Model | 16.599 | 0.001 | 0.354 | 0.1765 | 0.1281 | 24.847 | 18.040 | 0.001 | 0.001 | 0.001 | - | - |

**Table S6.** Collinearity between enviromental variables used in db-RDAs to examine the biogeographical determinants of DNA methylation in Canada lynx.

| Variance Inflation Factor | | |
|---|---|---|
| Distance | Insularity | Climate |
| 2.0932 | 2.747 | 3.7715 |

**Table S7.** Results of a gene ontology analysis, identifying biological functions of differentially methylated regions between Newfoundland and mainland Canada lynx.

| Biological Process | Class ID | Functional Hits | Genes | Expected | Fold Enriched | +/- | *p-value* |
|---|---|---|---|---|---|---|---|
| embryo development | GO:0009790 | 3 | LMO4, LRRC7, DLG1 | 0.19 | 15.67 | + | 0.000994 |
| → developmental process | GO:0032502 | 7 | NRG3, LMO4, LRRC7, DLG1, PPP3CA, PDZD2, TMOD2 | 2.71 | 2.58 | + | 0.0168 |
| visual perception | GO:0007601 | 2 | PDE3A, PDZD2 | 0.14 | 14.2 | + | 0.00915 |
| → system process | GO:0003008 | 5 | LRRC7, DLG1, PDE3A,PDZD2, TMOD2 | 1.84 | 2.71 | + | 0.0357 |
| → single-multicellular organism process | GO:0044707 | 7 | CDH18, NRG3, LRRC7, DLG1, PDE3A, PDZD2, TMOD2 | 3.01 | 2.33 | + | 0.028 |
| → multicellular organismal process | GO:0032501 | 7 | CDH18, NRG3, LRRC7, DLG1, PDE3A, PDZD2, TMOD2 | 3.04 | 2.3 | + | 0.0295 |
| negative regulation of apoptotic process | GO:0043066 | 2 | LMO4, HDAC9 | 0.18 | 11.19 | + | 0.0143 |
| protein glycosylation | GO:0006486 | 2 | CHSY3 | 0.19 | 10.55 | + | 0.0159 |
| cellular component morphogenesis | GO:0032989 | 4 | CDH18, LMO4, TMOD2, UTRN | 0.76 | 5.24 | + | 0.00711 |
| → cellular component organization | GO:0016043 | 8 | CDH18, LMO4, LRRC7, HDAC9, PRKCE, DLG1, TMOD2, UTRN | 3.55 | 2.26 | + | 0.0222 |
| → cellular component organization or biogenesis | GO:0071840 | 8 | CDH18, LMO4, LRRC7, HDAC9, PRKCE, DLG1, TMOD2, UTRN | 3.79 | 2.11 | + | 0.0498 |
| Unclassified | | 11 | GALNTL6, ZEB1, MTUS2, DCC, FAM210A, AUTS2, FAM35A, ANK2, CNTN5, GRIP1, CADM2 | 18.43 | 0.6 | - | 0.0217 |

**Table S8.** Methylated adapters used for multiplexing double-digest RRBS libraries, as well as the overall design, from van Gurp *et al.* (van Gurp et al. 2016). X indicates 5mC.

| **Barcode** | **AseI (Top)** | **AseI (Bottom)** |
|---|---|---|
| AACT | 5'-AXAXTXTTTXXXTAXAXGAXGXTXTTXXGATXTNNNaaxtC-3' | 5'-TAGagttNNNAGTTAGATCGGAAGAGCGTCGTGTAGGGAAAGAGTGT-3' |
| CCTA | 5'-AXAXTXTTTXXXTAXAXGAXGXTXTTXXGATXTNNNxxtaC-3' | 5'-TAGtaggNNNAGTTAGATCGGAAGAGCGTCGTGTAGGGAAAGAGTGT-3' |
| TTAC | 5'-AXAXTXTTTXXXTAXAXGAXGXTXTTXXGATXTNNNttaxC-3' | 5'-TAGgtaaNNNAGTTAGATCGGAAGAGCGTCGTGTAGGGAAAGAGTGT-3' |
| AGGC | 5'-AXAXTXTTTXXXTAXAXGAXGXTXTTXXGATXTNNNaggxC-3' | 5'-TAGgcctNNNAGTTAGATCGGAAGAGCGTCGTGTAGGGAAAGAGTGT-3' |
| GAAGA | 5'-AXAXTXTTTXXXTAXAXGAXGXTXTTXXGATXTNNNgaagaC-3' | 5'-TAGtcttcNNNAGTTAGATCGGAAGAGCGTCGTGTAGGGAAAGAGTGT-3' |
| CCTTC | 5'-AXAXTXTTTXXXTAXAXGAXGXTXTTXXGATXTNNNxxttxC-3' | 5'-TAGgaaggNNNAGTTAGATCGGAAGAGCGTCGTGTAGGGAAAGAGTGT-3' |
| TTCAA | 5'-AXAXTXTTTXXXTAXAXGAXGXTXTTXXGATXTNNNttxaaC-3' | 5'-TAGttgaaNNNAGTTAGATCGGAAGAGCGTCGTGTAGGGAAAGAGTGT-3' |
| GCGGC | 5'-AXAXTXTTTXXXTAXAXGAXGXTXTTXXGATXTNNNgxggxC-3' | 5'-TAGgccgcNNNAGTTAGATCGGAAGAGCGTCGTGTAGGGAAAGAGTGT-3' |
| AGATGC | 5'-AXAXTXTTTXXXTAXAXGAXGXTXTTXXGATXTNNNagatgxC-3' | 5'-TAGgcatctNNNAGTTAGATCGGAAGAGCGTCGTGTAGGGAAAGAGTGT-3' |
| CATAGC | 5'-AXAXTXTTTXXXTAXAXGAXGXTXTTXXGATXTNNNxatagxC-3' | 5'-TAGgctatgNNNAGTTAGATCGGAAGAGCGTCGTGTAGGGAAAGAGTGT-3' |
| TTCGAC | 5'-AXAXTXTTTXXXTAXAXGAXGXTXTTXXGATXTNNNttxgaxC-3' | 5'-TAGgtcgaaNNNAGTTAGATCGGAAGAGCGTCGTGTAGGGAAAGAGTGT-3' |

| | | |
|---|---|---|
| ATGCGC | 5'-AXAXTXTTTXXXTAXAXGAXGXTXTTXXGATXTNNNatgxgxC-3' | 5'-<br>TAGgcgcatNNNAGTTAGATCGGAAGAGCGTCGTGTAGGGAAAGAGTGT-3' |
| | **NsiI (Top)** | **NsiI (Bottom)** |
| AACT | 5'-GagttNNNAGATCGGAAGAGCGGTTCAGCAGGAATGCCGAG-3' | 5'-XTXGGXATTXXTGXTGAAXXGXTXTTXXGATXTNNNaaxtCTGXA-3' |
| CCAG | 5'-GctggNNNAGATCGGAAGAGCGGTTCAGCAGGAATGCCGAG-3' | 5'-XTXGGXATTXXTGXTGAAXXGXTXTTXXGATXTNNNxxagCTGXA-3' |
| TTGA | 5'-GtcaaNNNAGATCGGAAGAGCGGTTCAGCAGGAATGCCGAG-3' | 5'-XTXGGXATTXXTGXTGAAXXGXTXTTXXGATXTNNNttgaCTGXA-3' |
| GGTC | 5'-GgaccNNNAGATCGGAAGAGCGGTTCAGCAGGAATGCCGAG-3' | 5'-XTXGGXATTXXTGXTGAAXXGXTXTTXXGATXTNNNggtxCTGXA-3' |
| ACTA | 5'-GtagtNNNAGATCGGAAGAGCGGTTCAGCAGGAATGCCGAG-3' | 5'-XTXGGXATTXXTGXTGAAXXGXTXTTXXGATXTNNNaxtaCTGXA-3' |
| CAGC | 5'-GgctgNNNAGATCGGAAGAGCGGTTCAGCAGGAATGCCGAG-3' | 5'-XTXGGXATTXXTGXTGAAXXGXTXTTXXGATXTNNNxagxCTGXA-3' |
| TGAT | 5'-GatcaNNNAGATCGGAAGAGCGGTTCAGCAGGAATGCCGAG-3' | 5'-XTXGGXATTXXTGXTGAAXXGXTXTTXXGATXTNNNtgatCTGXA-3' |
| GTCG | 5'-GcgacNNNAGATCGGAAGAGCGGTTCAGCAGGAATGCCGAG-3' | 5'-XTXGGXATTXXTGXTGAAXXGXTXTTXXGATXTNNNgtxgCTGXA-3' |

5'-AXAXTXTTTXXXTAXAXGAXGXTXTTXXGATXTNNNBBBBC-3'
3'-TGTGAGAAAGGGATGTGCTGCGAGAAGGCTAGANNNBBBBGAT-5'

AseI

5'- GBBBBNNNAGATCGGAAGAGCGGTTCAGCAGGAATGCCGAG-3'
3'-AXGTCBBBBNNNTXTAGXXTTXTXGXXAAGTXGTXXTTAXGGXTX-5'

NsiI

**Table S9.** Total reads and cytosines identified for the non-methylated lambda phage DNA control. Conversion rate was calculated by dividing the methylated cytosines by the total number of cytosines analyzed.

| Total Reads | Methylated Cytosines | Non-Methylated Cytosines | Conversion Rate |
|---|---|---|---|
| 4,260,069 | 16,374,124 | 173,911,904 | 0.91395 |

**Table S10.** Total cytosines retained for analyses after filtering for shared positions between 95 individuals, subdivded by the feature-based sensitivity analysis. Unique positions are listed first, with total analyzed cytosines in parentheses.

| Methylation Dataset | Total Cytosines Analyzed | Total 5KB Windows | Average Coverage |
|---|---|---|---|
| Unannotated Regions | 5,031 (67,279) | 376 | 43.00 ±16.89% |
| CpG Islands / Gene Bodies | 4,611 (58,305) | 329 | 38.23 ± 23.92% |

**Table S11.** Illumina primers used during PCR. The forward primer is the Illumina PE PCR Primer 1.0, while the reverse is the PE PCR Primer 2.0.

| Illumina Primers | |
|---|---|
| Forward Primer | 5’- AATGATACGGCGACCACCGAGATCTACACTCTTTCCCTACACGACGCTCTTCCGATCT-3’ |
| Reverse Primer | 5’- TCTAGCCTTCTCGCCAAGTCGTCCTTACGGCTCTGGCTAGAGCATACGGCAGAAGACGAAC-3’ |

**Table S12.** Cycling conditions for PCR with bisulfite-converted DNA, using an uracil sensitive polymerase.

| PCR Conditions (20X) | | |
|---|---|---|
| 95°C | | 3 Minutes |
| | 98°C | 10 Seconds |
| | 65°C | 15 Seconds |
| | 72°C | 15 Seconds |
| 72°C | | 5 Minutes |

**Table S13**. Pairwise FST between Canada lynx populations at putatively neutral SNPs (n = 85).

| | AK | MB | QC | NL |
|---|---|---|---|---|
| AK | - | | | |
| MB | 0.1 | - | | |
| QC | 0.1 | 0.09 | - | |
| NL | 0.17 | 0.16 | 0.1 | - |

**References for SI reference citations**


1. van Gurp TP, et al. (2016) epiGBS: reference-free reduced representation bisulfite sequencing. *Nat Methods* 13(4):322–324.
2. Guo W, et al. (2017) CGmapTools improves the precision of heterozygous SNV calls and supports allele-specific methylation detection and visualization in bisulfite-sequencing data. *Bioinformatics* 34(February):381–387.
3. Lischer HEL, Excoffier L (2012) PGDSpider: An automated data conversion tool for connecting population genetics and genomics programs. *Bioinformatics* 28(2):298–299.
4. Foll M, Gaggiotti O (2008) A genome-scan method to identify selected loci appropriate for both dominant and codominant markers: A Bayesian perspective. *Genetics* 180(2):977–993.
5. Danecek P, et al. (2011) The variant call format and VCFtools. *Bioinformatics* 27(15):2156–2158.
6. Gutenkunst RN, Hernandez RD, Williamson SH, Bustamante CD (2009) Inferring the Joint Demographic History of Multiple Populations from Multidimensional SNP Frequency Data. *PLoS Genet* 5(10):e1000695.
7. Krueger F, Andrews SR (2011) Bismark: A flexible aligner and methylation caller for Bisulfite-Seq applications. *Bioinformatics* 27(11):1571–1572.
8. Laine VN, et al. (2016) Evolutionary signals of selection on cognition from the great tit genome and methylome. *Nat Commun* 7:1–9.
9. Maechler M, Rousseeuw P, Struyf A, Hubert M, Hornik K (2018) cluster: Cluster Analysis Basics and Extensions. R package version 2.0.7-1.
10. Jombart T (2008) Adegenet: A R package for the multivariate analysis of genetic markers.

*Bioinformatics* 24(11):1403–1405.

11. Brown J, Bennet J, French C (2017) SDMtoolbox 2.0: the next generation Python-based GIS toolkit for landscape genetic, biogeographic and species distribution model analyses. *PeerJ* 5(7):694–700.

12. Fick SE, Hijmans RJ (2017) WorldClim 2: new 1-km spatial resolution climate surfaces for global land areas. *Int J Climatol* 37(12):4302–4315.

13. DeFries RS, Hansen MC, Townshend JRG, Janetos AC, Loveland TR (2000) A new global 1-km dataset of percentage tree cover derived from remote sensing. *Glob Chang Biol* 6(2):247–254.

14. R Core Team (2017) R: A language and environment for statistical computing. R Foundation for Statistical Computing, Vienna, Austria. URL https://www.R-project.org/. Online.

15. Dixon P (2003) VEGAN, a package of R functions for community ecology. *J Veg Sci* 14(6):927–930.

16. Barratt CD, et al. (2018) Vanishing refuge? Testing the forest refuge hypothesis in coastal East Africa using genome-wide sequence data for seven amphibians. *Mol Ecol* (February):4289–4308.

17. Portik DM, et al. (2017) Evaluating mechanisms of diversification in a Guineo-Congolian tropical forest frog using demographic model selection. *Mol Ecol* 26(19):5245–5263.
